# Evolutionary analyses of base-pairing interactions in DNA and RNA secondary structures

**DOI:** 10.1101/419341

**Authors:** Michael Golden, Ben Murrell, Oliver G. Pybus, Darren Martin, Jotun Hein

## Abstract

Pairs of nucleotides within functional nucleic acid secondary structures often display evidence of coevolution that is consistent with the maintenance of base-pairing. Here we introduce a sequence evolution model, MESSI, that infers coevolution associated with base-paired sites in DNA or RNA sequence alignments. MESSI can estimate coevolution whilst accounting for an unknown secondary structure. MESSI can also use GPU parallelism to increase computational speed. We used MESSI to infer coevolution associated with GC, AU (AT in DNA), GU (GT in DNA) pairs in non-coding RNA alignments, and in single-stranded RNA and DNA virus alignments. Estimates of GU pair coevolution were found to be higher at base-paired sites in single-stranded RNA viruses and non-coding RNAs than estimates of GT pair coevolution in single-stranded DNA viruses, suggesting that GT pairs do not stabilise DNA secondary structures to the same extent that GU pairs do in RNA. Additionally, MESSI estimates the degrees of coevolution at individual base-paired sites in an alignment. These estimates were computed for a SHAPE-MaP-determined HIV-1 NL4-3 RNA secondary structure and two corresponding alignments. We found that estimates of coevolution were more strongly correlated with experimentally-determined SHAPE-MaP pairing scores than three non-evolutionary measures of base-pairing covariation. To assist researchers in prioritising substructures with potential functionality, MESSI automatically ranks substructures by degrees of coevolution at base-paired sites within them. Such a ranking was created for an HIV-1 subtype B alignment, revealing an excess of top-ranking substructures that have been previously identified as having structure-related functional importance, amongst several uncharacterised top-ranking substructures.

## 1 Introduction

The primary role of nucleic acid molecules, such as DNA (deoxyribonucleic acid) and RNA (ribonucleic acid), is to encode genetic information for storage and transfer. However, both types of molecules can form structures with additional functions (Mattick, 2003). DNA is ordinarily thought of as a double-stranded molecule forming the now iconic double helical configuration (Watson and Crick, 1953), although many viral genomes consist entirely of single-stranded DNA (ssDNA) or single-stranded RNA (ssRNA) molecules. Such single-stranded nucleic acid molecules are far less constrained than double-stranded ones in the variety of functional structures that they can form. For example, the Rev response element (RRE) within the single-stranded HIV RNA genome plays a crucial role in the regulation of HIV replication by binding the HIV Rev protein to facilitate the transfer of HIV genomes from the nucleus to the cytoplasm where translation and virion packaging occur (Heaphy *et al.*, 1990; Daugherty *et al.*, 2010).

The structures that nucleic acid molecules form are commonly referred to as their secondary or tertiary structures. Secondary structure is defined as the set of hydrogen bonding interactions between the constituent bases of a nucleic acid molecule; tertiary structure is defined as the arrangement of the constituent atoms of a nucleic acid molecule in three-dimensional space. This study focuses exclusively on RNA and DNA secondary structures.

Both computational (Markham and Zuker, 2008; Sükösd *et al.*, 2012; Bernhart *et al.*, 2006) and hybrid experimental-computational techniques (Wilkinson *et al.*, 2006) for secondary structure prediction exist. However, even if the secondary structure of an RNA sequence can be accurately determined, this does not immediately say anything about the potential functional or biological importance of the identified structure. Many RNA secondary structures are known to have specific biological functions, and it is expected that evolutionary conservation or adaptation of these structures might detectably impact patterns of sequence diversity and evolution.

One evolutionary signal that can be used to identify selectively maintained secondary structures is nucleotide coevolution. Nucleotide coevolution is expected at base-paired nucleotide positions within RNA and DNA secondary structures (Eddy and Durbin, 1994; Tuplin *et al.*, 2002; Cheng *et al.*, 2012). Many pairs of nucleotides within RNA molecules exhibit evidence of coevolution, such that whenever a substitution occurs in one partner of the pair, complementary substitutions are selected for in the other partner in a manner that is consistent with the selective maintenance of canonical base-pairing (Cheng *et al.*, 2012). The restricted nature of base-pairing interactions in nucleic acid structures (compared to amino acid interactions in protein structures) permits both nucleic acid structural conformations and nucleotide coevolution to predicted with relative ease. In this study we consider the canonical RNA base-pairs to be the two Watson-Crick base-pairs, GC and AU, and the weaker GU wobble base-pair (GC, AT, and GT base-pairs in DNA, respectively).

Methods for detecting coevolution, such as mutual information (Eddy and Durbin, 1994; Lindgreen *et al.*, 2006), can be used to aid the computational inference of secondary structures. Accordingly, some RNA comparative secondary structure prediction approaches, such as PPfold (Sükösd *et al.*, 2012), use information about coevolving nucleotides inferred from sequence alignments to more accurately predict secondary structures. Conversely, within a given secondary structural element, evidence that paired bases are coevolving is evidence of the functional importance of that element (Tuplin *et al.*, 2002; Cheng *et al.*, 2012; Muhire *et al.*, 2014).

Standard approaches for measuring coevolution (or more accurately: covariation), such as mutual information, are non-evolutionary in that they do not take into account the phylogenetic relationships of the sequences being analysed. Founder substitutions can, by chance, induce correlations between bases in a large number of observed variants or species (for an example see: Bhattacharya *et al.* (2007)), which may be mistaken for strong evidence of coevolution if the phylogeny is not accounted for. Substitution models provide a probabilistic framework for modelling of both phylogenetic relationships and underlying substitution processes.

In this article, we introduce MESSI (Modelling the Evolution of Secondary Structure Interactions), a probabilistic model that generalises upon the pioneering Muse (1995) (M95) model of base-pairing evolution. The first way in we extend the M95 model is the addition of parameters that allow us to differentiate between rates of evolution affecting the three canonical base-pairs. We used this to compare the role of GU base-pairs in single-stranded RNA viruses with GT base-pairs in single-stranded DNA viruses.

It is well-established that GU pairs can hydrogen bond in RNA to form base-pairs, although they are chemically weaker than GC and AU base-pairings (Rousset *et al.*, 1991). The relative chemical strengths of GC, AU, and GU base-pairs are partially due to the number of hydrogen bonds that form between their constituent bases: three for GC base-pairs, two for AU base-pairs, and two for GU base-pairs. Although GU pairs form the same number of hydrogen bonds as in AU pairs, the geometry of the bases leads to the GU pairing being substantially weaker than the AU pairing (Varani and McClain, 2000). Despite the weaker chemical interaction, GU base-pairings are known to be involved in functional RNA structures (Gautheret *et al.*, 1995). Less well understood is the role of GT base-pairings in DNA. There are few reports of GT base-pairings in double-stranded DNA helices (Early *et al.*, 1978; Ho *et al.*, 1985). Whilst, we were unable to directly measure the chemical strength of these base-pairing interactions in the present study, we used MESSI to analyse alignments for evidence of evolutionary forces favouring GT pairs at base-paired positions.

The second way in which we extended the M95 model was to allow substitution rates across to vary across sites (Yang, 1993, 1994), including allowing the two positions involved in a base-pairing to each to have a potentially different substitution rate. This was done to account for site-specific substitution rates, such as those expected within coding sequences. This is particularly important for virus genomes, where the majority of nucleotides are in protein coding regions, where some of these nucleotides additionally participate in functionally important base-pairing interactions.

The third extension was permit the strength of co-evolution to vary across base-paired sites. This provides a measure of base-pairing coevolution between every pair of sites in alignment, allowing us to test whether a particular pair of sites is coevolving in a manner favouring canonical base-pairing, or whether the two sites are evolving independently of one another. The use of an evolutionary model addresses the problem of founder effects potentially inflating signals of covariation. We used this extension to estimate rates of coevolution at individual base-paired sites within two HIV alignments, allowing us to identify and rank substructures within the larger HIV genomic secondary structure that have potential biological functionality. This is a feature of our model that we expect will assist researchers in focusing their experimental analyses on those portions of large RNA or DNA secondary structures that are most likely to be biologically relevant.

Compared to non-evolutionary methods, the computational cost of applying evolutionary models, such as MESSI, can severely limit their utility. We used GPU (graphics processing unit) parallelism and a Metropolis-within-Gibbs procedure when performing Bayesian inference to reduce these computational bottlenecks. This provided large speed-ups. Furthermore, this allowed us to account for a potentially unknown secondary structure configuration, whilst simultaneously estimating parameters of interest. This implies that the user need not provide a secondary structure as input. Relying on a potentially incorrect input secondary structure may bias parameter estimates, and may also undermine the conclusions of hypothesis tests based on those estimates. A further benefit of accounting for an unknown secondary structure is that this enables MESSI to output a prediction of the secondary structure and a base-pairing probability matrix.

## 2 Methods

### 2.1 The Muse 1995 model

Muse (1995) developed a paired site model, henceforth referred to as the M95 model. M95 accounts for RNA base-pairing constraints by modelling pairs of nucleotide positions using a 16 × 16 matrix. The model generalises upon standard 4×4 nucleotide substitution models, such as the GTR model, by introducing a coevolution parameter, *λ*, that is intended to capture substitutions at paired positions that are consistent with the maintenance of canonical RNA base-pairing. We define the set of canonical base-pairs as follows:

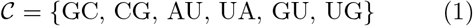

Equation 2 presents a version of the original M95 paired model based on a GTR model *Q* and a set of canonical base-pairs *𝒞*:

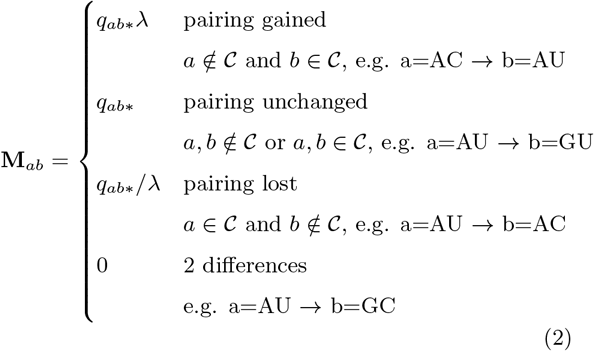

Where *a* and *b* are nucleotide pairs, *q*_*ab\**_ is the entry of the GTR matrix *Q* corresponding to the nucleotide position within the nucleotide pair that underwent a substitution, and *λ* is a parameter capturing the degree of RNA coevolution; *i.e.* the degree to which canonical RNA base-pairing is evolutionary maintained (*λ* > 1) or disrupted (*λ* < 1). Note that *λ* = 1 represents the *neutral* case, in which each of the two nucleotide positions in a pair are treated as evolving independently under the GTR model specified by *Q*.

Furthermore, let *π*^dinuc^ denote a length 16 vector of paired frequencies. *π*^dinuc^ is the union of two mutually exclusive cases: *π*^dinuc^ = *π*^unpaired^ ∪ *π*^paired^, *π*^unpaired^ represents the cases where the target pair *d*_*ij*_ is not in the set of canonical base-pairs (*d*_*ij*_ ∉ *𝒞*), and *π*^paired^ represents the cases where the target pair *d*_*ij*_ is in the set of canonical base-pairs (*d*_*ij*_ ∈ *𝒞*), respectively:

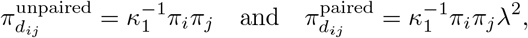

Note that *i* and *j* correspond to the first and second positions of the target pair, respectively. Where *π*_*i*_ is the equilibrium frequency under the GTR model, Q, of the nucleotide in the first position of the target pair *d*_*ij*_, and similarly *π*_*j*_ is the equilibrium frequency of the nucleotide in the second position. *κ*_1_ = 1 + 2(*π*_G_*π*_C_ + *π*_A_*π*_U_ + *π*_G_*π*_U_)(*λ*^2^ − 1) is a normalising constant that ensures the entries of *π*^*d*^ sum to one.

Note that within the set of canonical base-pairs, *𝒞* (defined in Equation 1), there are three pairs of symmetrical base-pairs: {GC, CG}, {AU, UA}, and {GU, UG}. It is assumed that each base-pair within a symmetrical pair has the same fitness. This is a reasonable assumption as it treats the evolution of nucleotides towards the 5’ end of the sequence the same as nucleotides towards the 3’ end. From this point forward we assume this symmetry and refer to the three pairs of symmetrical base-pairs as the *three canonical base-pairs*.

In the formulation of the original M95 model in Equation 2 all three canonical base-pairs in the set *𝒞* are treated as having equal fitness. However, there is good evidence that GU base-pairings in RNA, for example, are deleterious evolutionary intermediates relative to GC and AU (Rousset *et al.*, 1991). In light of this, in the next section we extend the M95 model such that substitutions affecting the three canonical base-pairs are not constrained to have the same rate of coevolution.

### 2.2 Differentiating between types of base-pairing substitutions

We extend the M95 model to differentiate between the three different canonical base-pairs, by introducing potentially distinct coevolution rates (*λ*_GC_, *λ*_AU_, and *λ*_GU_) for each of three different base-pairs (GC, AU, and GU, respectively). Using similar notation as in Equation 2, the extended rate matrix is given as follows:

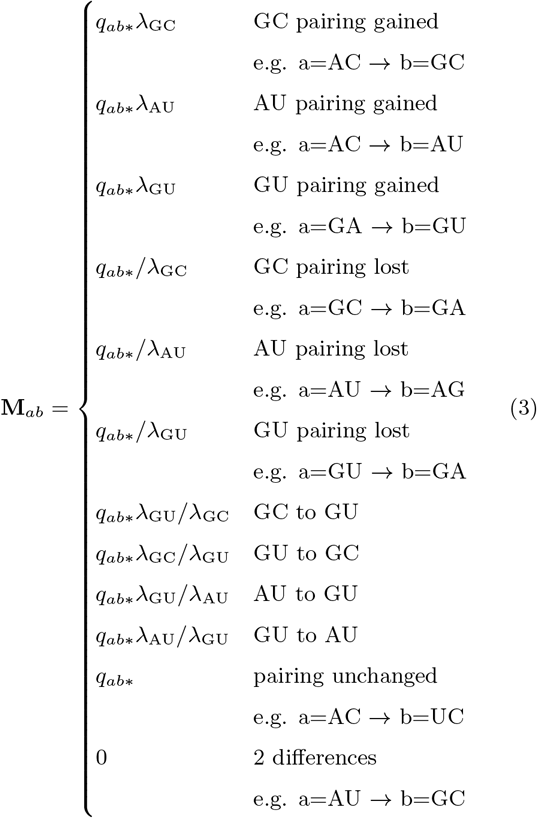

and the corresponding paired frequencies are:

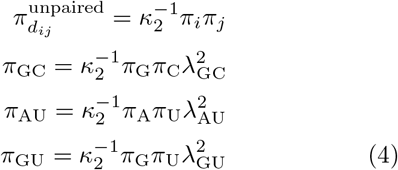

Where 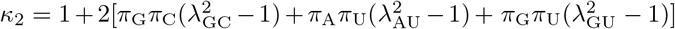

### 2.3 Stationarity and time-reversibility

We are able show for the extended model that the paired frequencies, *π*, given in (4) correspond to the stationary distribution of **M** by verifying that:

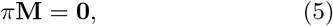

and that time-reversibility of **M** holds:

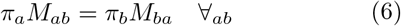

where *a* and *b* represent nucleotide pairs. The conditions in (5) and (6) were verified using the symbolic math package, SymPy (Joyner *et al.*, 2012), as implemented in the `musesymbolic.py` script (see Supplementary Material).

### 2.4 Modelling variable degrees of co-evolution

In the M95 model (2) the rate of coevolution was assumed to be the same for each base-paired site within a secondary structure 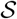. However, it is expected that the strength of the selective forces maintaining canonical base-pairing will vary amongst base-paired sites in 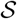. In this section, we extend the M95 model such that the *degree of coevolution*, denoted by *η*_*q,r*_, is able to vary from base-paired site to base-paired site. *η*_*q,r*_ is drawn independently for each base-paired site (described in the next section), and acts to scale the three coevolution rates as follows:

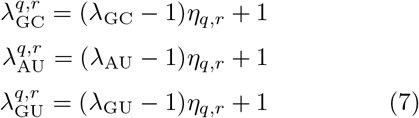

where *λ*_GC_ ≥ 1, *λ*_AU_ ≥ 1, and *λ*_GU_ ≥ 1 are the base-pairing substitution rates shared across all paired sites. This parametrisation was chosen so that 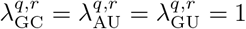 when *η*_*q,r*_ = 0.

In addition to allowing the rate of coevolution, *η*, to vary across base-paired sites, we also allow substitution rates to vary from site to site following the gamma distributed sites rate approach of (Yang, 1993, 1994). For unpaired sites, sequence evolution is modelled using a standard GTR+Γ model. For base-paired sites slightly more care needs to be taken (see Supplementary Section 1.2 for details). We call the version of our generalised M95 model that differentiates between the three canonical base-pairs and takes into account site-to-site rate variation, the ‘unconstrained M95 model’.

### 2.5 Testing neutrality of coevolution

To test the hypothesis that two nucleotide positions within a particular base-paired site are evolving neutrally, i.e. the substitutions at each of the two sites are occuring independently rather than actively favouring the maintenance canonical base-pairing, we assume that the degree of coevolution, *η*_*q,r*_, at each base-paired site is distributed as follows: *η*_*q,r*_ = 0 with probability *w*_*η*_ (the neutral, independent case), otherwise with probability 1 − *w*_*η*_, *η*_*q,r*_ is drawn from a discretised gamma distribution with M categories (the dependent case). Note that *η*_*q,r*_ ≥ 0 and therefore the case where substitutions are acting to disrupt canonical RNA base-pairing is not considered, i.e. the case where the coevolution parameters are between 0 and 1. For all analyses a discretisation of *M* = 4 was used, resulting in five rate categories: one neutral category with probability *w*_*η*_, and four positive categories each with probability 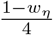.

### 2.6 Parameters

Table 1 lists the parameters and their distributions used in the most general version of the implemented model (the unconstrained model). Note that for some analyses we perform Bayesian inference, whereas for others we perform maximum likelihood (ML) inference. The distributions over the parameters specified here are those used for Bayesian inference, however, we also indicate how the parameters are treated during ML inference. Parameters are either estimated whilst ignoring the prior distribution, or fully marginalised under the prior distribution. Note that the phylogenetic tree, 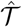, relating the alignment of sequences, *𝓓*, for both Bayesian and ML inference is estimated in advance and fixed *a priori* using Fast-Tree (Price *et al.*, 2010) under a GTR+CAT model.

**Table 1:** Parameters of the unconstrained model and their distributions.

| Parameter and distribution | Marginalised or estimated | Description |
| --- | --- | --- |
| $w_\eta \sim \text{Beta}(2,2)$ | Estimated | Probability of neutral coevolution. |
| $X_{q,r} \sim \text{Bernoulli}(w_\eta)$ | Marginalised | Indicates neutral coevolution at position $q, r$ when $X_{q,r} = 1$. |
| $c \sim \text{Exponential}(\frac{1}{10})$ | Estimated | Shape and rate parameter of prior over coevolution rates $\eta_{q,r}$. |
| $\eta_{q,r} = 0$ if $X_{q,r} = 1$, otherwise:<br>$\eta_{q,r} \sim \text{DiscretisedGamma}_M(c, c)$ | Marginalised | The rate of coevolution at each paired position $q, r$. |
| $d \sim \text{Exponential}(\frac{1}{10})$ | Estimated | Shape and rate parameter of prior over substitutions rates $\rho_q$. |
| $\rho_q \sim \text{DiscretisedGamma}_K(d, d)$ | Marginalised | Substitution rate at each unpaired position $q$. |
| $(\pi_A, \pi_C, \pi_G, \pi_T) \sim \text{Dir}(1, 1, 1, 1)$ | Estimated | GTR equilibrium frequencies of the four nucleotides. |
| $q_{AC} \sim \text{Exponential}(\frac{1}{10})$ | Estimated | GTR rate matrix entry AC. |
| $q_{AG} \sim \text{Exponential}(\frac{1}{10})$ | Estimated | GTR rate matrix entry AG. |
| $q_{AT} \sim \text{Exponential}(\frac{1}{10})$ | Estimated | GTR rate matrix entry AT. |
| $q_{CG} \sim \text{Exponential}(\frac{1}{10})$ | Estimated | GTR rate matrix entry CG. |
| $q_{CT} \sim \text{Exponential}(\frac{1}{10})$ | Estimated | GTR rate matrix entry CT. |
| $q_{GT} \sim \text{Exponential}(\frac{1}{10})$ | Estimated | GTR rate matrix entry GT. |
| $\lambda_{GC} \sim \text{Exponential}(\frac{1}{10}) + 1$ | Estimated | GC coevolution rate. |
| $\lambda_{AT} \sim \text{Exponential}(\frac{1}{10}) + 1$ | Estimated | AT coevolution rate. |
| $\lambda_{GT} \sim \text{Exponential}(\frac{1}{10}) + 1$ | Estimated | GT coevolution rate. |
| $\mathcal{S} \sim \text{KH99}$ | Marginalised | The secondary structure is drawn from the KH99 SCFG prior. |

### 2.7 Computer representations of secondary structure

To model nucleic acid secondary structure a suitable definition of secondary structure is required. We use the definition outlined in Moulton *et al.* (2000): a secondary structure, 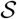, for a nucleic acid molecule consisting of *N* nucleotides is a simple graph specified by the vertex set [*N*]:= {1, …, *N*} and an edge set 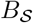. Where each vertex in [*N*] corresponds to a nucleotide and each edge in the edge set 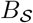 corresponds to a base-pair. 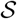 is such that if 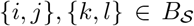 with *i* < *j* and *k* < *l* then:

i. *i* = *k* if and only if *j* = *l*, and
ii. *k* ≤ *j* implies that *i* < *k* < *l* < *j*

Vertices that are not contained within the edge set 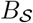 are termed *unpaired*. Condition (i) implies that each vertex (nucleotide) belongs to at most one base-pair. Condition (ii) prevents *pseudoknotting*, *i.e.* non-nested base-pairs.

Note that pseudoknotting is physically possible in both real RNA and DNA structures, but is excluded in many definitions of secondary structures as efficient algorithms exist for marginalising or maximising over secondary structures when assuming (ii). Our method permits a canonical secondary structure with pseudoknots to be specified *a priori*, however, if the user instead treats the structure as unknown, MESSI will strictly marginalise over non-pseudoknotted structures only.

Figure 1 gives a computational format for representing secondary structures. The dot-bracket format (Figure 1A) is a natural and compact way of representing non-pseudoknotted secondary structures. Matching brackets represent base-paired nucleotide positions and dots represent unpaired (singled-stranded) nucleotide positions. To represent pseudoknotted structures (structures that violate condition (*ii*)) additional bracket types are required (Figure 1D).

**Figure 1:**
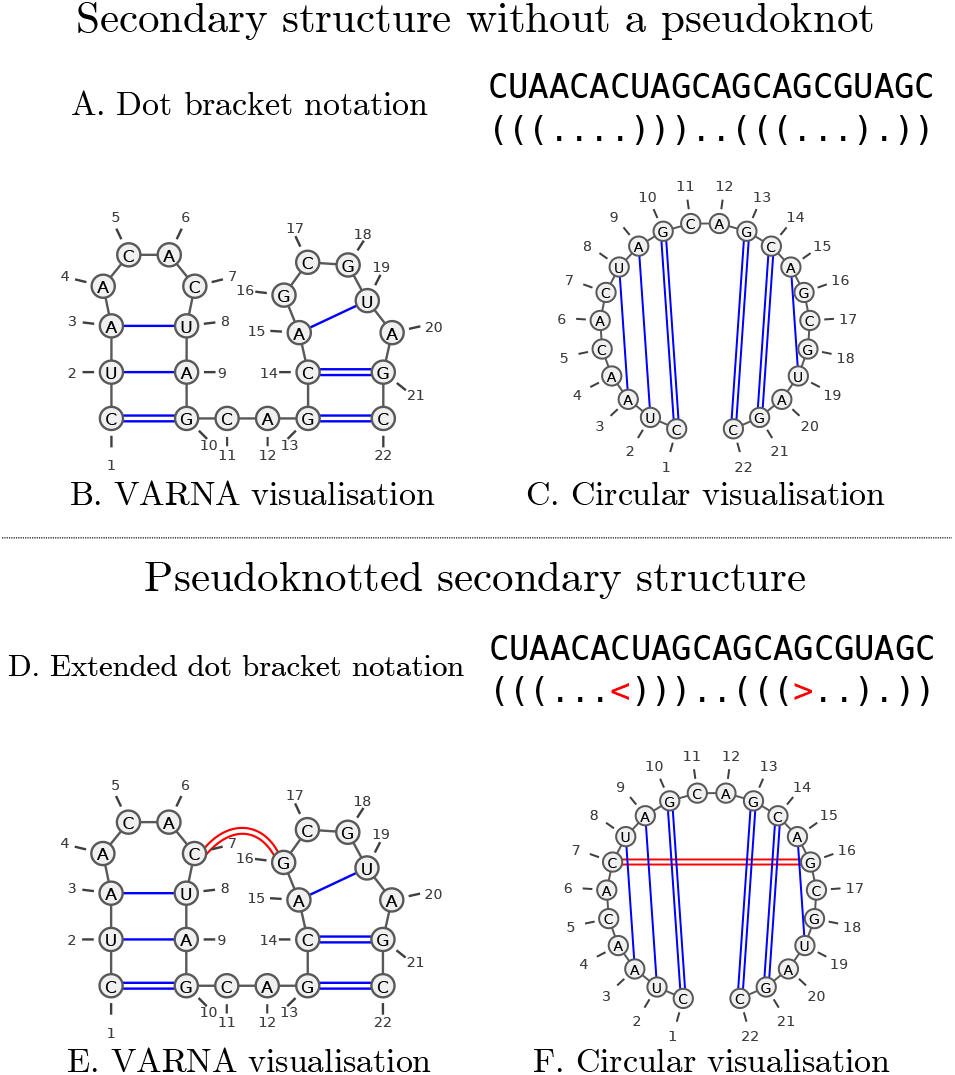
Examples of secondary structure representations. Above (A) is a dot bracket representation of a secondary structure, and the corresponding VARNA and circular visualisations (B and C, respectively) produced by VARNA Darty *et al.* (2009). Below (D) is an extended dot bracket notation format with an additional bracket type, *<>*, that allows a pseudoknotted structure to be represented unambiguously. E and F are the corresponding VARNA visualisations for D. Note how the overlapping bonds in the circular visualisation (F) demonstrate that the secondary structure is pseudoknotted.

### 2.8 Likelihood

Conditioned on a secondary structure, 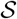, unpaired nucleotide positions within 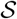, denoted by 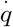, and base-paired nucleotide positions within 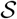, denoted by 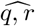, are assumed to be independent. The likelihood of an alignment, *𝓓*, is given by a simple product of unpaired and paired site likelihoods:

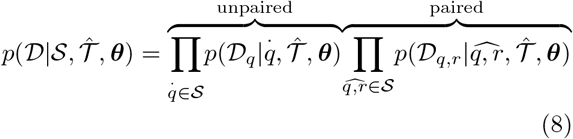

where 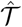 is a phylogenetic tree. Felsenstein’s pruning algorithm (Felsenstein, 1981) was used to calculate both the unpaired site likelihoods, 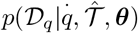, and the paired site likelihoods,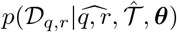. Paired sites were modelled using the unconstrained M95 model, whereas unpaired sites were modelled using the GTR + Γ model that is nested within the unconstrained M95 model.

### 2.9 Prior over RNA secondary structures

Equation 8 assumes that the secondary structure 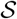 is known *a priori*, either through experimental or computational methods of structure prediction. However, it also possible to treat the secondary structure as unknown, by placing a prior probability distribution, 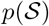, over secondary structures and marginalising 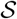.

One way of introducing a prior over secondary structures is by using a Stochastic Context Free Grammar (SCFG). A SCFG is probabilistic extension of a context-free grammar (CFG). A CFG is a type of grammar that defines a set of rules for generating all possible strings in a given formal language. A SCFG extends this notion by assigning probabilities to each possible string in the given language. RNA SCFGs are SCFGs that give probability distributions over strings of base-paired and unpaired nucleotides representing RNA secondary structures (Anderson, 2014).

#### 2.9.1 The KH99 grammar

We chose the KH99 SCFG (Knudsen and Hein, 1999) as a prior over secondary structures. The rules and associated probabilities for this SCFG are given as follows:

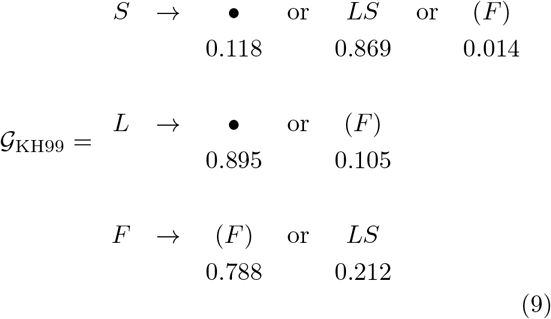

Note that *S* is the start symbol.

The KH99 assigns probabilities to all strings of a specified length that can be written in dot-bracket notation, with at least two unpaired nucleotides separating every base-pair.

#### 2.9.2 Structure-integrated likelihood

Using Bayes’ rule, the probability of a secondary structure, 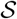, conditional on the data, *𝓓*, and phylogenetic parameters, ***θ***, is given by:

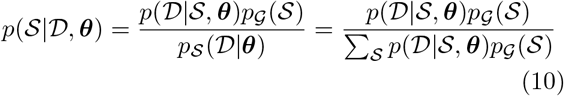

We take particular note of the *structure-integrated likelihood* term in the denominator of (10):

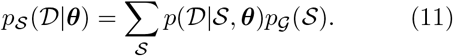

This term requires summing over all possible secondary structures and is not a constant that can be ignored due it’s dependence on ***θ***. This number grows exponentially with the length of the alignment *L*. Fortunately, there exists an *𝓞*(*L*^3^) polynomial-time algorithm, the inside algorithm (Lari and Young, 1991), for summing the probabilities of all derivations of an SCFG (all valid secondary structures in the case of RNA SCFG). By analogy to the forward algorithm for HMMs, the inside algorithm allows the structure-integrated likelihood, 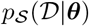 (the analogue of the forward likelihood for HMMs), to be efficiently computed. The structure-integrated likelihood is given by element *I*(*S*, 1, *L*) of the inside probability matrix, where *S* is the start symbol of the KH99 grammar.

Likewise, by analogy to the backward algorithm for HMMs, there exists an ‘outside algorithm’, which together with the inside probabilities allows the posterior marginals of the hidden variables to be computed (in the case of an RNA SCFG, these are the positional emission probabilities of base-pair and unpaired terminal symbols – see Supplementary Section 1.5).

#### 2.9.3 Parallelisation of the inside and outside algorithms

Algorithm’s S1 and S2 in the Supplementary Material provide pseudocode for iterative implementations of the inside and outside algorithms for SCFGs in double emission normal form, respectively.

Figure 2 illustrates the calculation of the inside probability matrix, showing the order in which elements are computed and the data dependencies required to compute a particular element. Using these patterns, Sükösd *et al.* (2011) developed a strategy for CPU parallelism, whereby blocks of elements running diagonally along the inside matrix can be computed in parallel, as they do not have data dependencies. We implemented a similar scheme for the CUDA GPU architecture, whereby instead of blocks, each element along a diagonal is computed in parallel. This can be done because each element along a diagonal is independent of all other elements on the same diagonal. For large alignments (*L* > 1000) this implies thousands of computational threads executing the same set of instructions in parallel, but on different data (different elements of a particular diagonal), this is known as SIMD (Single Instruction Multiple Data) parallelism and is the regime of parallelism for which GPU architectures are tailored. As far as we are aware this is the first GPU implementation of the inside and outside algorithms.

**Figure 2:**
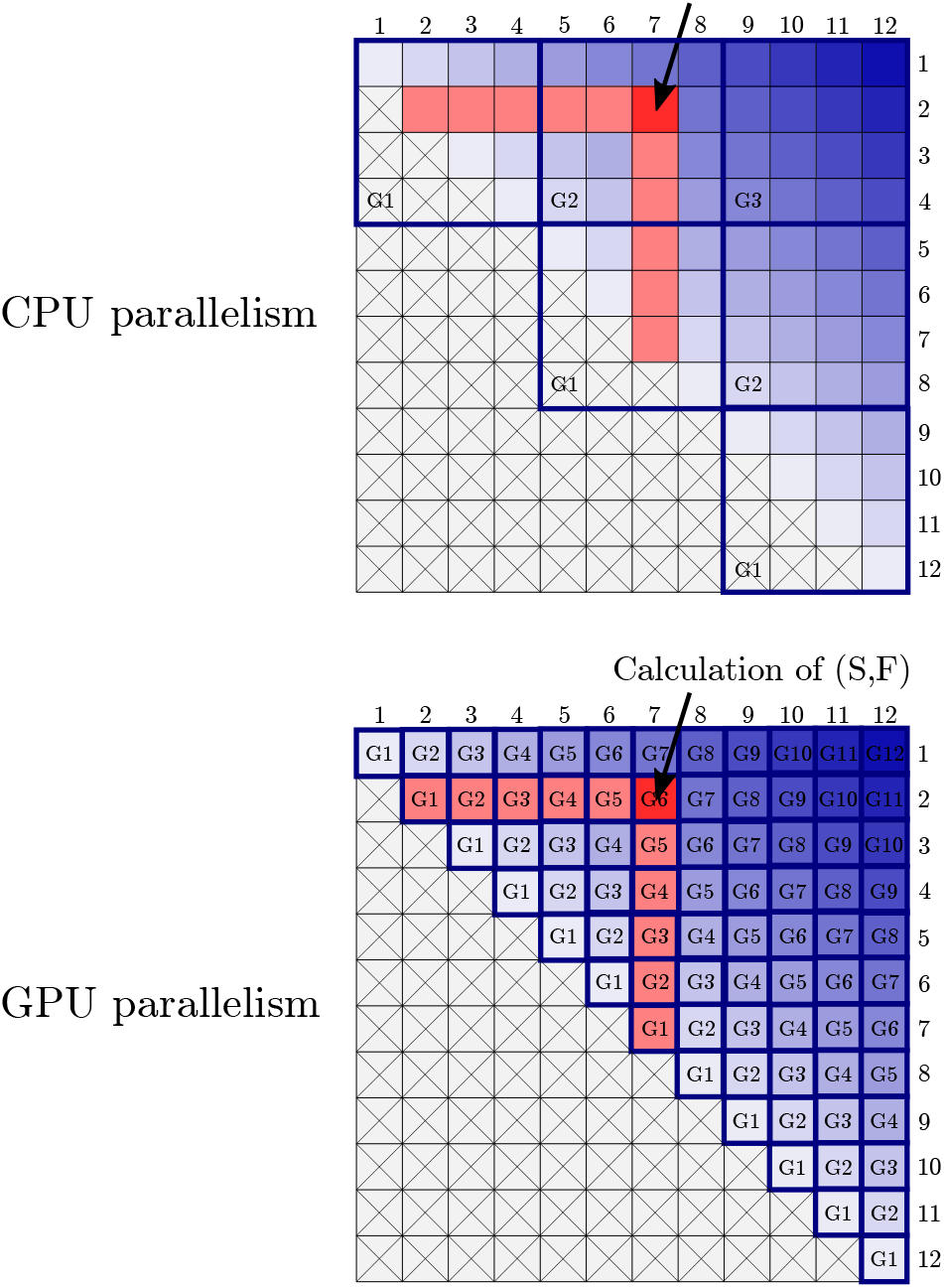
Illustrations of the inside algorithm showing CPU and GPU parallelism schemes. The light to dark blue gradient starting at the central diagonal and finishing in the top right-hand corner indicates the order in which each diagonal is computed. The light red elements indicate the data dependencies required to compute the single bright red entry of the inside matrix. The lower half of each matrix with each cell crossed out is not computed and can be ignored. Note that the top-right element corresponds to the structure-integrated likelihood term and is therefore always the last element to be calculated, as it depends on all other elements having been computed first.

### 2.10 Paired site likelihoods

Because the inside and outside algorithms consider every possible base-pairing they require a matrix **B** of *paired site likelihoods*. Each element **B**_*qr*_ of **B** corresponds to a paired site likelihood 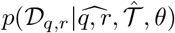 for a pair of sites, *q* and *r*, in the alignment *𝓓*, which can be calculated using Felsenstein’s peeling algorithm. Since the diagonal of **B** is ignored and **B**_*qr*_ = **B**_*rq*_ (i.e. **B** is symmetric), 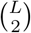 paired site likelihoods need to be calculated. Whilst, the number of computational steps is only *𝓞*(*L*^2^) in the alignment length *L*, compared to *𝓞*(*L*^3^) for the inside and outside algorithms, the amount of time per computational step for computing the paired site likelihoods is substantially higher due the use of Felsenstein’s algorithm. To ameliorate this bottleneck, we use the partial site caching strategy of Pond and Muse (2004) to reduce the number of likelihood calculations required and developed a CUDA GPU implementation.

Note that the inside and outside algorithms also require a vector, **S**, of length of *L* single site likelihoods, where each element corresponds to 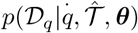. However, this is fast to compute compared to the matrix **B**.

### 2.11 Sampling secondary structure configurations

The inside probability matrix can be used to sample secondary structure configurations from the distribution

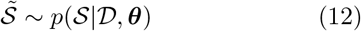

Sampling terminal strings (secondary structures in our case) using an SCFG is analogous to sampling hidden state sequences using the forward-filtering backward-sampling algorithm for HMMs (Frühwirth-Schnatter, 1994). An algorithmic description for sampling secondary structures from an RNA SCFG is given in the Supplementary Methods Section 1.4.

### 2.12 Bayesian posterior inference

The posterior distribution of the continuous-parameters, ***θ***, conditional on the data *𝓓* and a secondary structure 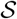 can be sampled using the Metropolis-Hastings algorithm and the relationship given by Bayes’ formula:

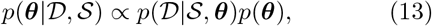

where the likelihood term, 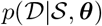, is given by (8) and *p*(***θ***) is the prior.

We can also treat the secondary structure as unknown and assume a RNA SCFG prior,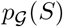, over secondary structures. This can be achieved by using the structure-integrated likelihood, 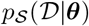, when inferring ***θ***:

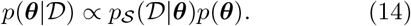

However, note that the structure-integrated likelihood term is computed every time a new set of parameters is proposed. As mentioned previously, this requires computing a matrix **B** of paired site likelihoods (requiring *𝓞*(*L*^2^) computational steps) and calculating the final structure-integrated likelihood term using the inside algorithm (requiring *𝓞*(*L*^3^) computational steps). Therefore gathering enough samples to ensure an adequate sample size will be relatively slow. However, given that we can sample the conditional distribution, 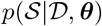, using the sampling procedure outlined in Section 2.11 this leads to a potentially more efficient Metropolis-within-Gibbs approach. This approach works by alternatively sampling from the full conditional distribution:

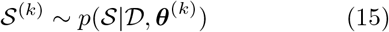

using the sampling procedure outlined in Section 2.11 and

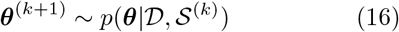

using the Metropolis-Hastings algorithm. Whilst the Gibbs sampling step (15) still requires computing a matrix **B** of paired site likelihoods and running the inside algorithm, the Metropolis-Hastings step (16) only requires *𝓞*(*L*) operations and can be repeated for multiple iterations following the Gibbs sampling step. In our implementation we repeat the Metropolis-Hastings step 50 times following the Gibbs sampling step. Together these give a Markov Chain Monte Carlo algorithm whose stationary distribution, 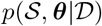, and associated marginals, 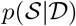 and *p*(***θ***|**𝓓**), are the distributions of interest.

### 2.13 Maximum likelihood inference

The COBYLA optimization algorithm (Powell, 1994) in the NLOpt library (Johnson, 2014) was used to find the maximum likelihood (ML) parameters via the structure-integrated likelihood (11). Note that when doing so the priors over the continuous parameters were either ignored and estimated using ML, or the priors were used and the parameters were fully marginalised (as specified in the *Priors* section).

### 2.14 Site permutations

To test whether secondary structure dependencies present in real datasets influence model fit, each alignment was taken and its sites randomly permuted. Two such nucleotide column permuted datasets (*p*_1_ and *p*_2_) were generated for each real dataset. ML estimation using the structure-integrated likelihood was used to fit the parameters of each permuted dataset under the unconstrained model and the secondary structure information entropy was calculated.

## 3 Results

### 3.1 Site permutation benchmarks

To assess the degree to which secondary structure dependencies present in real datasets influence model fit, ML inference was performed on real and permuted datasets, and their structure-integrated likelihoods and structure information entropies were compared (Section 2.14 in Methods). The structure-integrated likelihoods for the permuted datasets were expected to be lower than those of the real datasets. Note that comparing these likelihoods is valid given they are in effect marginal likelihoods. Conversely, the structure information entropies were expected to be higher for the permuted datasets than for the real datasets. Unlike the real datasets, the patterns of coevolution in the permuted datasets were not expected to coincide with stable secondary structure configurations, thereby spreading the probability mass over a larger number of secondary structure configurations.

The maximum likelihood estimates of the structure-integrated likelihoods were indeed lower for the permuted datasets in every instance (Supplementary Table S1). This partially validates our model and is consistent with the presence of real secondary structure dependencies in the original datasets. As expected, the structure information entropies were higher for the permuted datasets, with the exception of RF00003, which had marginally lower structure information entropies for both of the permuted datasets. This result is surprising as RF00003 corresponds to the U1 spliceosomal RNA, a component of a spliceosome (Burge *et al.*, 2012) with a thermodynamically stable structure. Since MESSI uses evolutionary and not thermodynamic information to infer secondary structure, one explanation may be that the patterns of nucleotides within the RF00003 dataset are only weakly informative of the underlying secondary structure.

### 3.2 Benchmarks of RNA structure prediction

Whilst, our model was not designed to predict RNA secondary structure, the expected base-pairing and unpairing probabilities can be calculated (see Supplementary Section 1.5) and a Maximum Expected Accuracy consensus secondary structure determined (see Supplementary Section 1.6). Our method was compared to two comparative methods of RNA secondary structure prediction: RNAalifold (Bernhart *et al.*, 2006) and PPfold (Sükösd *et al.*, 2012). The three methods were benchmarked on 22 alignments each having a corresponding experimentally-determined canonical RNA secondary structure from the RFAM database (Burge *et al.*, 2012). Five different measures were used to compute predictive accuracy (see Supplementary Section 1.8 for definitions of these measures).

MESSI has lower precision but higher recall than the other two methods, implying that it predicts more base-pairs (higher recall), but with a higher number of false-positives (lower precision; Figure 3). For the F1-score and MCC measures, both of which combine precision and recall, MESSI performs slightly better than RNAalifold and PPfold. MESSI also performs marginally better with respect to the mountain similarity measure – a measure that takes into account the overall ‘shape’ of the secondary structures being compared, rather than the exact matching of base-pairs.

**Figure 3:**
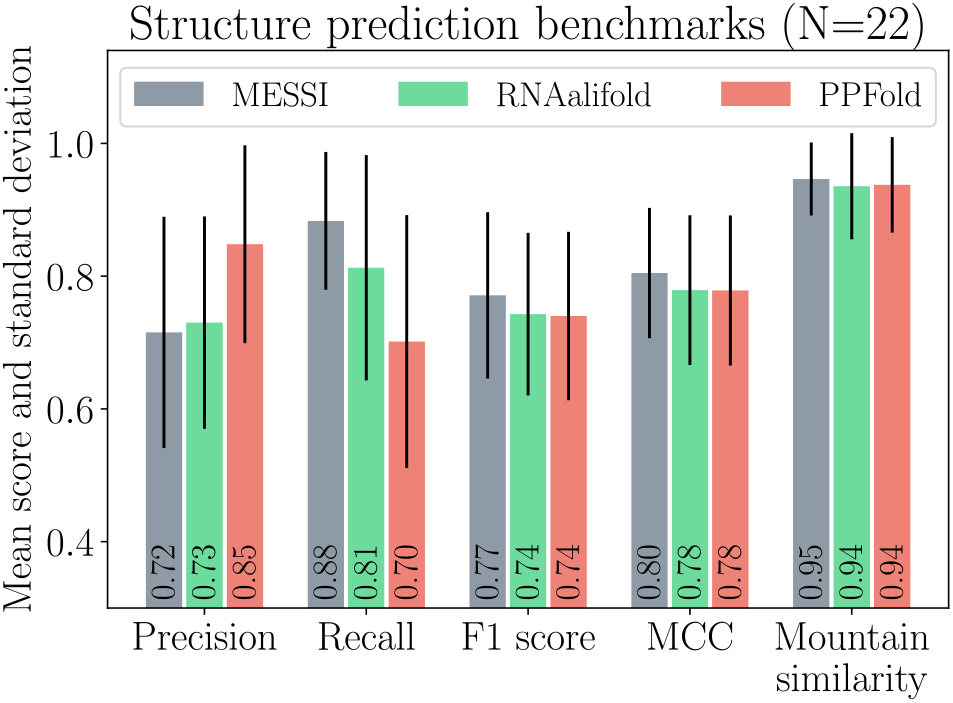
Summary of secondary structure prediction benchmarks. Structure predictions were performed on 22 RFAM datasets using three different comparative structure prediction methods (MESSI, RNAalifold, and PPFold).

Overall, our method performs on a par with two well-established methods of comparative RNA structure prediction. This was surprising given that the model was not developed for the purpose of secondary structure prediction. Maximum likelihood inference was used to estimate the model parameters. Where the coevolution parameters (*λ*_GC_, *λ*_AU_, and *λ*_GU_) were free to vary with the only restriction being: *λ*_GC_ ≥ 1, *λ*_AU_ ≥ 1, and *λ*_GU_ ≥ 1. Although not tested here, it might be possible to improve the predictive accuracy of MESSI’s structure predictions by performing Bayesian or MAP inference of the parameters using a set of priors whose hyperparameters are learnt from a training dataset of alignments and corresponding structures.

### 3.3 CPU and GPU timing benchmarks

The two computational bottlenecks in performing both maximum likelihood and Bayesian inference are computing paired site likelihood matrices (computed using an iterative version of Felsenstein’s algorithm) and computing inside probability matrices (using an iterative version of the inside algorithm); both of these steps are required repeatedly. Although optimised CPU implementations written in Julia were created for both of these steps, these were still relatively slow. Therefore GPU implementations written in CUDA were implemented for both.

The number of computational steps is expected to grow linearly with the number of unique paired site patterns and hence this was chosen as a predictor of the computational time required (Figure 4). Compared to the single-threaded CPU implementation we achieve a *∼* 50× speed-up with the GPU implementation across most datasets.

**Figure 4:**
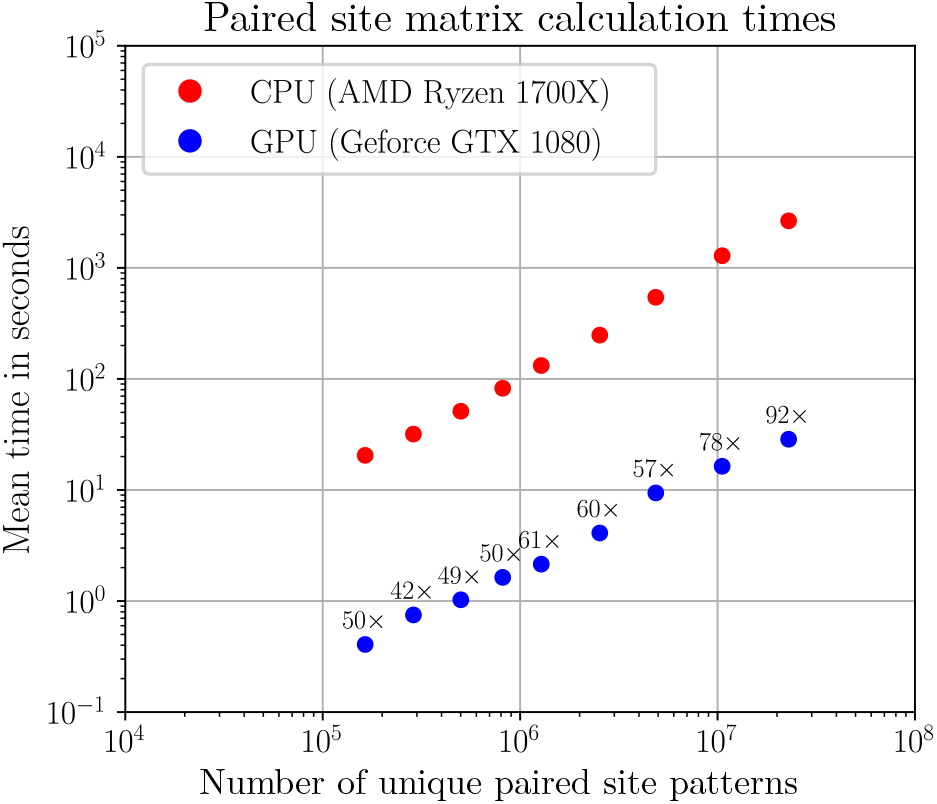
Paired site likelihoods calculation timings in seconds (log_10_ axis) as a function of the number unique paired partial site patterns (log_10_ axis). Numbers above the GPU timings indicate the fold speed-up over the CPU version.

The number of computational steps for the inside algorithm is expected to grow *𝓞*(*L*^3^) where *L* is the number of alignment sites in a particular dataset (Figure 5). A 50 to 200 fold speed-up for the paired site likelihood calculations was achieved for moderate dataset sizes, with the fold speed-ups for larger datasets being bigger, due to larger datasets better saturating the GPU.

**Figure 5:**
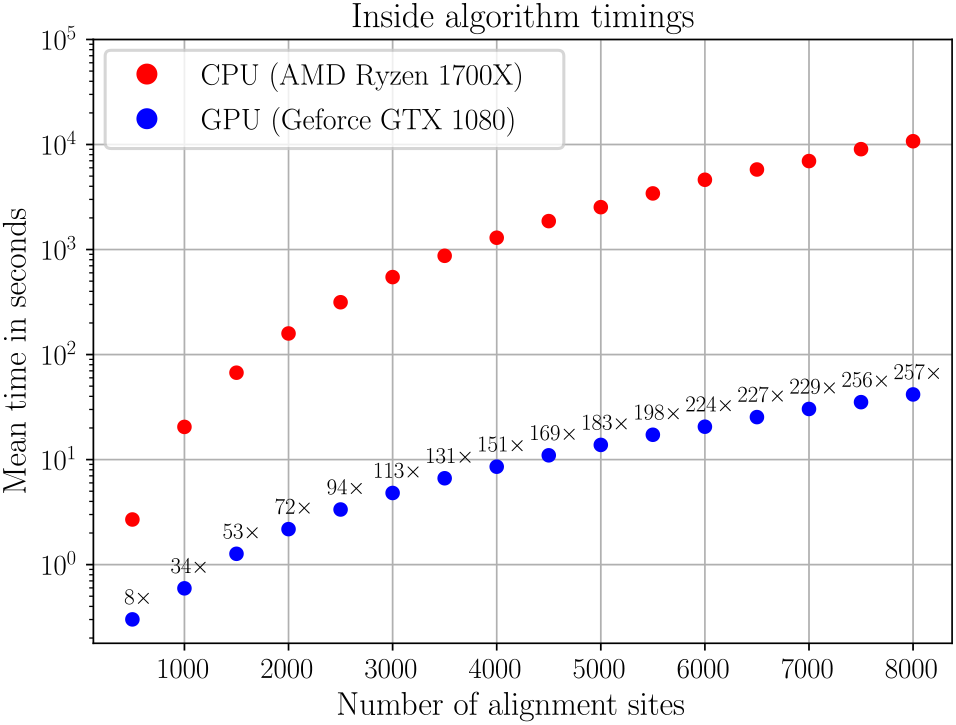
Inside algorithm timings in seconds (log_10_ axis) as a function of the number of alignment sites. Numbers above the GPU timings indicate the fold speed-up over the CPU version.

The speed-ups seen here are significant, enabling us to analyse datasets which would typically be considered intractable. Note that CPU and GPU implementations were also developed for the outside algorithm with similar speed-ups obtained (Figure S3 in the appendix).

### 3.4 The role of GU and GT base-pairs in single-stranded RNA and DNA

For all five non-coding RNA datasets (RF00001, RF00003, RF00010, RF00379, and RF01846) likelihood ratio tests (LRTs) rejected the GU neutral model in favour of the unconstrained model (*p* < 0.0005. See Table 2), with ML estimates for 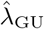 in the range 2.25 − 3.57. This is evidence that many GU pairs are under selective maintenance in the five non-coding RNA datasets tested.

**Table 2:** Tests of the GU/GT neutral hypothesis across 15 datasets: five non-coding RNA alignments from the RFAM database (denoted by the prefix ‘RF’), five ssRNA virus alignments (Foot-and-mouth-disease, Human poliovirus 1, Tobamovirus, Rhinovirus A, and Hepatitis A virus), and five ssDNA virus alignments (Maise Streak virus, Tomato Yellow Leaf Curl virus, Beet Curly Top virus, and Wheat Dwarf virus).

| Dataset | Type | Number of sites | GU (GT) neutral log-likelihood (M0) | Unconstrained log-likelihood (M1) | Delta M1-M0 log-likelihood | LRT p-value | $\hat{\lambda}_{GU}$ ($\hat{\lambda}_{GT}$) |
| --- | --- | --- | --- | --- | --- | --- | --- |
| RF01846 | ncRNA | 624 | -11101.49 | -11043.98 | 57.51 | *** | 2.64 |
| RF00379 | ncRNA | 335 | -10523.44 | -10470.32 | 53.12 | *** | 3.57 |
| RF00010 | ncRNA | 996 | -85067.98 | -84007.62 | 1060.36 | *** | 3.01 |
| RF00003 | ncRNA | 203 | -4711.20 | -4653.04 | 58.16 | *** | 2.97 |
| RF00001 | ncRNA | 230 | -28077.49 | -27897.66 | 179.83 | *** | 2.25 |
| FMDV | ssRNA | 8349 | -114986.98 | -114610.85 | 376.12 | *** | 5.21 |
| H. poliovirus 1 | ssRNA | 7668 | -65013.50 | -64966.32 | 47.19 | *** | 5.60 |
| Tobamovirus | ssRNA | 6849 | -88906.32 | -88767.09 | 139.23 | *** | 7.53 |
| Rhinovirus A | ssRNA | 7308 | -219614.83 | -217136.20 | 2478.62 | *** | 21.92 |
| Hepatitis A | ssRNA | 7572 | -63755.35 | -63755.31 | 0.04 | n.s. | 1.00 |
| MSV | ssDNA | 2755 | -15345.85 | -15341.48 | 4.37 | ** | 1.57 |
| TYLCV | ssDNA | 2925 | -40743.07 | -40743.03 | 0.04 | n.s. | 1.00 |
| BCTV | ssDNA | 3215 | -32094.09 | -32093.56 | 0.53 | n.s. | 1.18 |
| Bocavirus | ssDNA | 5577 | -31987.81 | -31985.98 | 1.84 | n.s. | 1.25 |
| WDV | ssDNA | 2755 | -10733.63 | -10731.59 | 2.04 | * | 1.57 |

\* $p < 0.05$ ; \*\* $p < 0.005$ ; \*\*\* $p < 0.0005$ ; n.s. = not significant

For four of the five RNA virus datasets tested (Rhinovirus A, Tobamovirus, human poliovirus 1, and foot-and-mouth disease virus. See Table 2) LRTs rejected the GU neutral model in favour of the unconstrained model (*p* < 0.0005 in all four cases), with ML estimates for 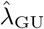 in the range 5.21 − 21.92. Curiously, the GU neutral model could not be rejected in favour of the unconstrained model for the hepatitis A virus dataset (Table 2), with the ML estimate for 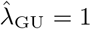

Three of the five DNA virus genome datasets tested (Human bocavirus, beet curly top virus, and tomato yellow leaf curl virus in Table 2) showed no significant difference between the unconstrained model and a GU (GT) neutral model (*λ*_GU_:= 1). In contrast, the Wheat Dwarf Virus dataset rejected the GT neutral model (*p* < 0.05), and the Maise Streak Virus dataset rejected the GT neutral model (*p <* 0.005). ML estimates for 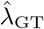 were in the range 1.0 − 1.57 for the five ssDNA virus datasets, which was low compared to those determined for the non-coding RNA and RNA virus datasets.

The LRT results and the ML estimates for 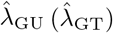 suggest that GT pairs are under weak selective maintenance in DNA virus genomes, and strong selective maintenance in RNA virus genomes and non-coding RNAs. This may indicate that GT base-pairings in DNA are chemically weaker relative to GU base-pairings in RNA and hence do not stabilise DNA secondary structures to the same extent as GU base-pairings in RNA.

### 3.5 Relative coevolution rates

The relative selective strengths of the coevolution rates associated with GC, AU and GU pairs were compared across both DNA and RNA virus genomes. The original M95 model assumed that *λ*_GC_:= *λ*_AU_ and *λ*_GC_:= 1. However, experimental evidence shows that GC base-pairings are chemically stronger than AU base-pairings in RNA (Mathews *et al.*, 1999), with both being substantially stronger than GU base-pairings.

To assess whether *λ*_GC_:= *λ*_AU_ is a reasonable assumption, we performed LRTs comparing the unconstrained model to a *λ*_GC_:= *λ*_AU_ constrained model. For 14 of the 15 datasets, LRTs rejected the GC-AU constrained model in favour of the unconstrained model (results not shown). The only exception was the human poliovirus 1 dataset, where the GC-AU constrained model could not be rejected.

This was explored further by comparing the inferred relative magnitudes of the rates associated with GC, AU (AT) and GU (GT) dinculeotides. If the fitness value of a RNA secondary structure element is positively correlated with its chemical stability, it is expected that the relative chemical stabilities associated with the three canonical base-pairs would be reflected in the relative magnitudes of the coevolution rates inferred by MESSI.

We applied MESSI’s Bayesian posterior inference mode to 15 datasets, five from each of three dataset types. Posterior probabilities associated with all six possible orderings of the three base coevolution rates were estimated for each dataset (Figure 6). Given the relative chemical base-pairing stabilities, the dominant ordering for the base coevolution rates was expected to be *λ*_GC_ > *λ*_AU_ > *λ*_GU_. For all five ncRNA datasets, all five ssDNA virus datasets, and two of the five ssRNA virus datasets the posterior probability associated with the *λ*_GC_ > *λ*_AU_ > *λ*_GU_ ordering was indeed 1.0.

**Figure 6:**
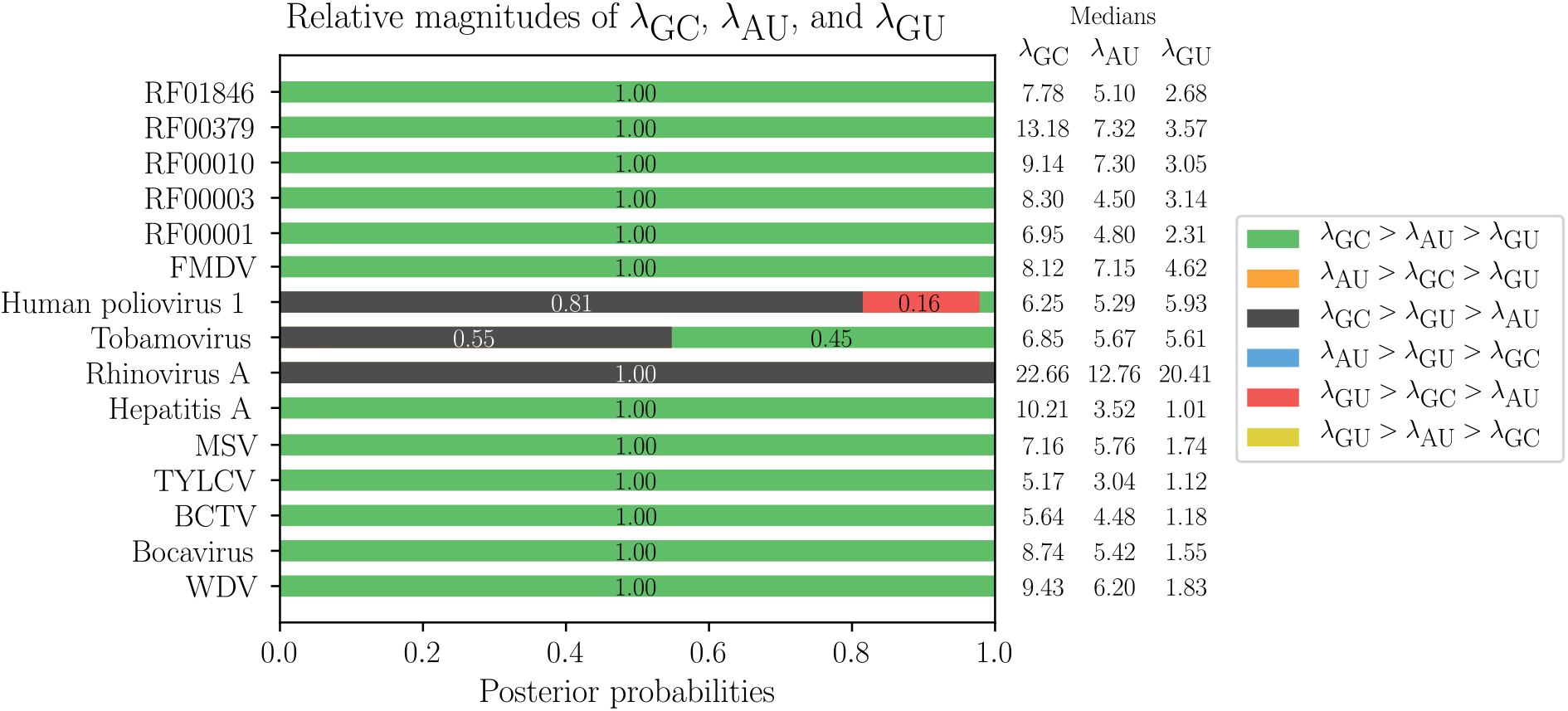
Estimated posterior probabilities for all six orderings of the three base coevolution rates across 15 datasets.

Interestingly, an unexpected ordering, *λ*_GC_ > *λ*_GU_ > *λ*_AU_, emerged for three of the ssRNA with a posterior probability of 1.0 for the Rhinovirus A dataset and posterior probabilities of 0.81 and 0.55 for the Human poliovirus 1 and Tobamovirus datasets, respectively. Possible explanations for this result include: (i) for many datasets it is not valid to assume a canonical secondary structure that is shared across the entire phylogeny, particularly in the presence of genome-scale ordered RNA structure (Simmonds *et al.*, 2004) which is not expected to be conserved. Two or more parts of the phylogeny may have different mutually exclusive secondary structures, giving rise to misleading patterns of pair evolution, and (ii) datasets with coding regions have additional constraints on synonymous and non-synonymous substitutions, and these protein coding constraints might mislead MESSI.

### 3.6 Degrees of coevolution are correlated with experimental SHAPE-MaP quantities

A notable example of a large RNA structure that has been partially experimentally-determined is that of the HIV-1M subtype B NL4-3 isolate (Watts *et al.*, 2009; Siegfried *et al.*, 2014). Rather than relying solely on computational techniques for the determination of RNA secondary structure of the 9173 nucleotide NL4-3 genome, the hybrid experimental-computational SHAPE-MaP (Selective 2’-hydroxyl acylation analysed by primer extension and mutational profiling; Siegfried *et al.* (2014)) approach was used to model the structure. The SHAPE-MaP approach preferentially mutates unpaired nucleotides, allowing the mutated nucleotides to be identified using DNA sequencing following reverse transcription. The SHAPE-MaP reactivity information is then used to constrain a thermodynamic RNA folding algorithm, enabling the construction of a secondary structure model which is reflective of the experimental data.

We compared three non-evolutionary computational measures of covariation (A. Mutual information, B. RNAalifold mutual information, and C. Mutual information with stacking; *Lindgreen et al.* (2006) and two evolutionary measures of coevolution inferred by MESSI (D. Posterior probability *η* ≠ 1, and E. Posterior mean *η*) with experimental SHAPE-MaP reactivities and SHAPE-MaP pairing probabilities at base-paired sites corresponding to three different datasets: an HIV 1b dataset, an HIV group 1M dataset, and a Simian Immunodeficiency Virus (SIV) dataset. When analysing the HIV datasets the SHAPE-MaP reactivities, SHAPE-MaP pairing probabilities and base-pairings were derived from a SHAPE-MaP analysis of the HIV NL4-3 sequence (Watts *et al.*, 2009). When analysing the SIV dataset a SHAPE-MaP analysis of the SIVmac239 sequence (Pollom *et al.*, 2013) was used. Given that high SHAPE-MaP reactivities indicate unpairing, we expected that degrees of co-evolution (or covariation) would be negatively correlated with SHAPE-MaP reactivities. Conversely, given that some paired nucleotides are expected to be selectively maintained due to structure-related functional importance, we expected a positive correlation between degrees of coevolution (or covariation) and SHAPE-MaP pairing probabilities.

For all three datasets the two measures of coevolution (D and E) were significantly correlated with both the SHAPE-MaP reactivities and SHAPE-MaP pairing probabilities using Spearman’s rank correlation test. The correlations were in the expected direction (negatively correlated for SHAPE-MaP reactivities and positively correlated for SHAPE-MaP pairing probabilities; Table 3). For all three datasets the correlation coefficients were significantly stronger in the expected direction for the two coevolution measures (D and E) than the the three covariation measures (A,B, and C; see the 95% confidences intervals for Spearman’s rho). We note that whilst many of the correlations were statistically significant, the magnitudes of the correlations were weak.

**Table 3:** Spearman’s correlations (*ρ*) and 95% confidence intervals (*ρ* 95% CI) between five different measures of covariation/coevolution and base-pair averaged SHAPE-MaP reactivities and the same five measures and base-pair averaged SHAPE-MaP pairing probabilities. Underlined values indicate correlations that are statistically significant and in the expected direction.

| Dataset | Measure | SHAPE-MaP reactivities |  |  | SHAPE-MaP pairing probabilities |  |  |
| --- | --- | --- | --- | --- | --- | --- | --- |
| | | $\rho$ | $\rho$ 95% CI | p-val | $\rho$ | $\rho$ 95% CI | p-val |
| HIV-1<br>subtype B | A. Mutual information (MI) | -0.01 | [-0.051, 0.035] | n.s. | <u>0.11</u> | [0.069, 0.154] | *** |
|  | B. RNAAlifold MI | 0.01 | [-0.033, 0.053] | n.s. | 0.01 | [-0.030, 0.056] | n.s. |
|  | C. MI with stacking | -0.02 | [-0.060, 0.026] | n.s. | <u>0.10</u> | [0.059, 0.144] | *** |
| | D. $p(\eta \neq 0)$ | <u>-0.16</u> | [-0.202, -0.118] | *** | <u>0.19</u> | [0.147, 0.230] | *** |
| | E. Posterior mean $\eta$ | <u>-0.14</u> | [-0.180, -0.095] | *** | <u>0.20</u> | [0.162, 0.244] | *** |
| HIV-1<br>group 1M | A. Mutual information (MI) | 0.03 | [-0.016, 0.070] | n.s. | 0.09 | [0.047, 0.132] | *** |
|  | B. RNAAlifold MI | 0.03 | [-0.013, 0.073] | n.s. | <u>0.08</u> | [0.033, 0.119] | *** |
|  | C. MI with stacking | 0.00 | [-0.043, 0.043] | n.s. | <u>0.12</u> | [0.081, 0.166] | *** |
| | D. $p(\eta \neq 0)$ | <u>-0.18</u> | [-0.225, -0.142] | *** | <u>0.27</u> | [0.227, 0.307] | *** |
| | E. Posterior mean $\eta$ | <u>-0.16</u> | [-0.197, -0.113] | *** | <u>0.29</u> | [0.251, 0.330] | *** |
| SIVmac239 | A. Mutual information (MI) | 0.09 | [0.053, 0.132] | *** | -0.04 | [-0.084, -0.005] | * |
|  | B. RNAAlifold MI | 0.12 | [0.086, 0.164] | *** | -0.07 | [-0.114, -0.035] | *** |
|  | C. MI with stacking | 0.10 | [0.057, 0.135] | *** | -0.01 | [-0.046, 0.033] | n.s. |
| | D. $p(\eta \neq 0)$ | <u>-0.12</u> | [-0.160, -0.082] | *** | <u>0.19</u> | [0.153, 0.229] | *** |
| | E. Posterior mean $\eta$ | <u>-0.10</u> | [-0.137, -0.058] | *** | <u>0.20</u> | [0.164, 0.240] | *** |

\* $p < 0.05$ ; \*\* $p < 0.005$ ; \*\*\* $p < 0.0005$ ; n.s. = not significant

Curiously, for the SIV dataset, SHAPE-MaP reactivities were significantly positively correlated with the three measures of covariation (A, B, and C) rather than negatively correlated as expected. There is broad evidence to suggest that base-paired sites in a functionally important RNA structure tend to be be more conserved (less variable) due to being under selective constraint (Muhire *et al.*, 2014; Tuplin *et al.*, 2004) and that double-stranded RNA (i.e. base-paired positions) is less susceptible to mutational processes (Lindahl and Nyberg, 1974). Conversely, unpaired sites are expected to undergo relatively higher rates of mutation. These higher rates of mutation may cause the three non-evolutionary measures of covariation to be erroneously inflated, given that they do not fully account for site-to-site rate variation (see Supplementary Section 1.2) unlike the coevolution measures inferred under our model, which do. It should also be noted that the SIV dataset is highly diverse compared to the two HIV datasets. Given these factors, it is anticipated that weakly base-paired sites will have inflated degrees of covariation using measures A, B and C, which may explain the unexpected positive correlation.

Overall, these results provide some reassurance that our method is performing as expected and that the evolutionary measures of coevolution are more reliable than the three measures of covariation that do not take into account evolutionary dependencies amongst the sequences being analysed. The detected degrees of coevolution suggest that a large proportion of the predicted base-pairings in the SHAPE-MaP structures have been selectively maintained since the common ancestors of the sequences being analysed in each of the three datasets.

### 3.7 Ranking and visualisation of sub-structures

Rather than considering the entire secondary structure of a large sequence, it is often useful to consider individual substructures. There are two primary reasons for considering substructures: (i) smaller regions are more easily conceptualised, and (ii) if functional components of a secondary structure are present, they tend to correspond to small regions (20-350 nucleotides long) of that secondary structure.

MESSI automatically ranks substructures by degrees of coevolution between their constituent nucleotides (see Supplementary Methods Section 1.9). We produced two rankings based on an HIV-1 subtype B alignment. The first ranking treated the HIV-1 NL4-3 SHAPE-MaP secondary structure as the canonical structure when inferring coevolution and identifying substructures (denoted the SHAPE structure ranking; Table 4 and Supplementary Table S3). The second ranking used a consensus structure estimated by MESSI based on base-pairing probabilities (denoted the consensus structure ranking; Table 5 and Supplementary Table S4).

**Table 4:** SHAPE structure ranking. The top 10 of 86 non-overlapping HIV NL4-3 substructures ranked from highest to lowest z-score based on the estimated degrees of coevolution within an alignment of HIV-1 subtype B sequences. Where the HIV NL4-3 SHAPE-MaP secondary structure was used as the canonical structure.

| Rank | Alignment position | NL4-3 position | Length | Name | Median degree of coevolution | z-score |
| --- | --- | --- | --- | --- | --- | --- |
| 1 | 8233 - 8582 | 7249 - 7595 | 350 | Rev Response element (RRE) | 5.38 | 5.02 |
| 2 | 2608 - 2943 | 1991 - 2326 | 336 | Longest continuous helix | 5.17 | 2.92 |
| 3 | 10155 - 10383 | 8982 - 9170 | 229 | 3' Untranslated region (3'UTR) | 5.27 | 2.69 |
| 4 | 588 - 838 | 105 - 344 | 251 | 5' Untranslated region (5'UTR) | 5.65 | 2.61 |
| 5 | 9570 - 9584 | 8440 - 8454 | 15 |  | 5.91 | 2.29 |
| 6 | 860 - 979 | 366 - 485 | 120 | 5' Untranslated region (5'UTR) | 5.54 | 2.28 |
| 7 | 1710 - 1845 | 1177 - 1312 | 136 |  | 5.17 | 2.28 |
| 8 | 2115 - 2301 | 1561 - 1711 | 187 | Gag-pol frameshift | 5.31 | 2.21 |
| 9 | 1479 - 1490 | 946 - 957 | 12 |  | 5.85 | 2.04 |
| 10 | 3886 - 3907 | 3269 - 3290 | 22 |  | 5.80 | 2.01 |

**Table 5:** Consensus structure ranking. The top 10 of 118 non-overlapping HIV consensus substructures ranked from highest to lowest z-score based on their degrees of coevolution within an alignment of HIV-1 subtype B sequences. Where the canonical structure was treated as unknown and a consensus structure predicted by MESSI.

| Rank | Alignment position | NL4-3 position | Length | Name | Median degree of coevolution | z-score |
| --- | --- | --- | --- | --- | --- | --- |
| 1 | 8240 - 8577 | 7256 - 7590 | 338 | Rev Response element (RRE) | 5.64 | 6.53 |
| 2 | 2202 - 2229 | 1645 - 1672 | 28 | Gag-pol frameshift | 8.17 | 4.56 |
| 3 | 1710 - 1845 | 1177 - 1312 | 136 |  | 6.44 | 4.50 |
| 4 | 4751 - 4833 | 4134 - 4216 | 83 |  | 6.47 | 3.97 |
| 5 | 4505 - 4709 | 3888 - 4092 | 205 |  | 5.22 | 3.21 |
| 6 | 591 - 939 | 108 - 445 | 349 | 5' Untranslated region (5'UTR) | 5.38 | 3.16 |
| 7 | 133 - 151 | NA | 19 |  | 6.85 | 2.94 |
| 8 | 2564 - 2890 | 1947 - 2273 | 327 | Longest continuous helix | 4.44 | 2.62 |
| 9 | 9782 - 9800 | 8645 - 8663 | 19 |  | 6.92 | 2.55 |
| 10 | 3612 - 3623 | 2995 - 3006 | 12 |  | 6.74 | 2.50 |

The highest ranked substructure in both the SHAPE and consensus rankings was the RRE (SHAPE RRE visualised in Figure 7). The RRE occurs in the genomes of all known HIV groups and plays a crucial role in the regulation of HIV virion expression (Heaphy *et al.*, 1990; Daugherty *et al.*, 2010).

The longest continuous helix identified in both the SHAPE-MaP and MESSI structures was ranked 2nd in the SHAPE ranking and 8th in the consensus ranking, respectively. The SHAPE-MaP analysis revealed that this helix is highly stable, although its function is unknown. The significant degrees of coevolution detected at base-paired sites within this substructure and the fact that MESSI detects it as conserved across all HIV-1 subtype sequences provides further evidence of its likely functional importance.

**Figure 7:**
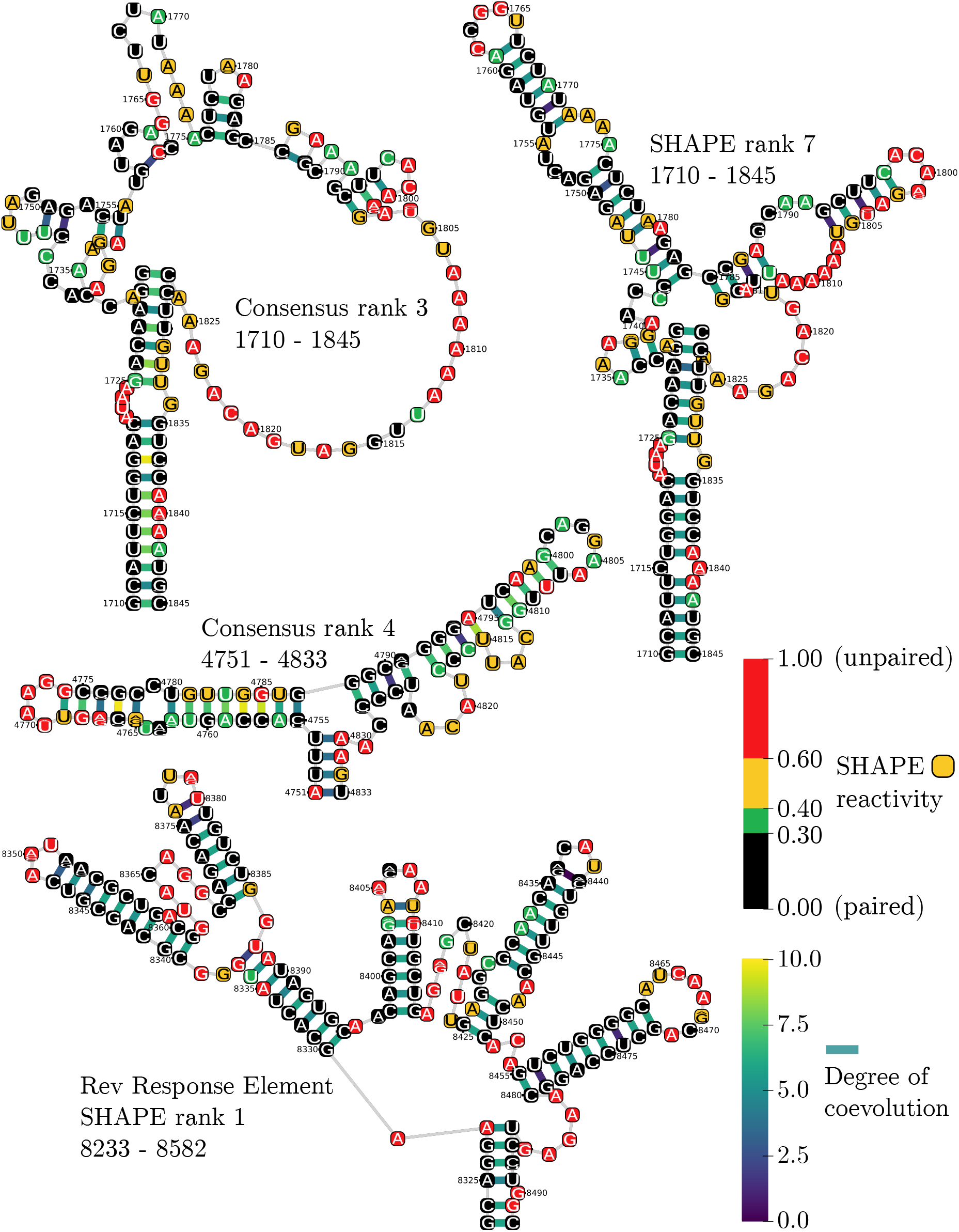
Visualisation of several top ranking substructures in the SHAPE-MaP structure and consensus structure rankings. NL4-3 SHAPE-MaP experimental reactivities are mapped and visually overlaid using the same colour scheme as in Watts *et al.* (2009). Depicted within each nucleotide is a sequence logo summarising the nucleotide composition at the corresponding alignment position. Mean degrees of coevolution inferred using MESSI are depicted for each base-pair using coloured links (blue-green-yellow gradient).

Portions of the 3’ and 5’ untranslated regions (UTRs) were ranked 3rd and 4th in the SHAPE ranking, respectively. This was not surprising given that these are both non-coding regions. The 5’ UTR is involved in regulation of translation (Damgaard *et al.*, 2004), whereas the 3’ UTR is believed to be involved in regulation of transcription (Watts *et al.*, 2009). A 5’ UTR substructure at a similar position is ranked 6th in the consensus ranking, whereas a 3’ UTR substructure at a similar position was not detected in the consensus structure. This may be explained by the large number of UTR missing sequences and high degrees of alignment uncertainty in the HIV-1 subtype B alignment in the UTR regions; factors which would both reduce support for the predicted base-pairings in the consensus structure.

An uncharacterised substructure (alignment position: 1710-1845) ranked 7th in the SHAPE structure ranking and 3rd in the consensus structure ranking (Figure 7). This substructure warrants further study, given the supporting evidence from experimental SHAPE-MaP reactivities, MESSI’s coevolution estimates, and MESSI’s evidence of conservation across HIV-1 subtype B sequences. Despite MESSI predicting the same helix as SHAPE-MaP at the base of this substructure, the remainder of the substructure is different in the SHAPE-MaP model. It is likely that the SHAPE-MaP model of this substructure is more accurate in this instance.

Interestingly, an additional uncharacterised substructure (alignment position: 4751-4833) ranked 4th in consensus ranking, but was not present in the HIV-1 NL4-3 SHAPE structure and hence was not present in the SHAPE structure ranking (Figure 7). Overlaid SHAPE-MaP reactivities from the HIV N4L-3 SHAPE model provide some support for MESSI’s prediction; particularly at unpaired positions which are supported by high SHAPE-MaP reactivities (indicating single-strandedness). It is possible that either MESSI’s or SHAPE-MaP’s prediction is wrong, or that the particular conformation predicted by MESSI is conserved amongst a subset of HIV-1 subtype B sequences that excludes NL4-3. It is also possible that this substructure exists in alternative conformations depending on *in vivo* or *in vitro* conditions.

Finally, the gag-pol frameshift-associated substructure was ranked 8th in the SHAPE ranking and 2nd in the consensus rankings. This substructure regulates the ratio of HIV gag/gag-pol that is expressed. Ribosomal synthesis of the gag-pol polyprotein requires a −1 ribosomal frameshift, without which translation ends in synthesis of the gag protein alone.

Overall, there is an excess of top-ranking substructures that have been identified previously in the literature as having structure-related importance. This is particularly evident in the SHAPE-MaP structure ranking. The use of the experimentally-determined SHAPE-MaP structure as the canonical structure strongly informs the SHAPE structure ranking, but has the disadvantage that it is based only on the HIV NL4-3 sequence rather than being representative of base-pairings conserved across all sequences within the HIV-1 subtype B alignment. By contrast, the consensus ranking canonical structure is predicted by MESSI and is based solely on evolutionary information, rather than experimental or thermodynamic information. In the future, we hope to extend MESSI by adding both experimental constraints from experiments such as SHAPE-MaP and thermodynamic constraints from folding software such as Vienna RNAfold. We expected this to improve estimates of coevolution and the overall ranking provided by MESSI.

## 4 Concluding remarks

MESSI was developed for modelling substitutions that are consistent with the maintenance of canonical base-pairing at paired sites within alignments of DNA and RNA sequences. To achieve this, we extended an existing model, M95 (Muse, 1995), in four major ways: (i) differentiating between the three canonical base-pairs (GC, AU and GU), (ii) allowing substitution rates to vary across sites, (iii) permitting the strength of coevolution to vary across base-paired sites, to measure the strength of selection operating on particualr base-pairs, and (iv) accounting for a potentially unknown secondary structure.

Amongst these extensions, extending the model to permit an unknown secondary structure posed the greatest computational challenges. The first challenge was the need to compute likelihoods using Felsenstein’s peeling algorithm for all 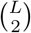 paired sites. Fortunately, a large number of redundant calculations could be avoided due to a large proportion of paired sites sharing the same partial site patterns (Pond and Muse, 2004), resulting in at least a 5× speed-up. Additionally, a further 50× speed-up was achieved using a GPU implementation of Felsenstein’s peeling algorithm. The second challenge was the need to marginalise an unknown secondary structure using the inside algorithm. Computational speed-ups of 50× - 200× were achieved using a GPU implementation of the inside algorithm. For Bayesian inference a Metropolis-within-Gibbs procedure was implemented to further avoid calculating the paired matrix likelihoods and inside probabilities at every iteration.

ML and Bayesian inference were used for different analyses. ML inference allowed us to perform likelihood ratio tests of various hypotheses, for which Bayesian model comparison was computationally intractable. Bayesian inference was used to obtain posterior distributions over various parameters, including the rates of coevolution associated with the three canonical base-pairs, and posterior probabilities and degrees of coevolution at base-pair sites.

To perform an initial validation of our model, site permutations of nucleotide alignments were performed to disrupt the secondary structure dependencies expected in real datasets. Consistent with the model behaving desirably, the structure-integrated maximum likelihood values were lower, and the structure information entropy values higher for the permuted datasets overall.

The ability to marginalise an unknown secondary structure shared amongst an alignment of sequences, implies that MESSI is also capable of secondary structure prediction. Although MESSI was not designed with structure prediction in mind, we found that it performed similarly to two popular comparative secondary structure prediction methods: RNAalifold (Hofacker, 2009) and PPfold (Sükösd *et al.*, 2012). This result further validates our approach.

We found strong evidence that GU pairs are selectively favoured at base-paired sites in five non-coding RNA datasets and four of five RNA virus genome datasets. Strong evidence for selection of GT pairs at base-paired sites was found for only one out of five of the DNA virus datasets tested. The notion that GU pairs play a role in stabilizing RNA secondary structures is consistent with numerous phylogenetic, and experimental analyses of RNA molecules (Woese *et al.*, 1980; Eddy and Durbin, 1994; Deigan *et al.*, 2009). The role of GT base-pairings in stabilizing DNA genomic secondary structures remains unclear.

We applied our model to the HIV-1 NL4-3 secondary structure and two corresponding alignment datasets containing large numbers of HIV-1 sequences, and an SIVmac239 secondary structure and a corresponding alignment of SIV sequences. We found that correlations between the SHAPE-MaP-determined quantities and degrees of coevolution as detected using MESSI were stronger than correlations between the same quantities and three non-evolutionary measures of covariation.

Interactive visualisations of the HIV-1 NL4-3 SHAPE-MaP and consensus secondary structures with the inferred degrees of coevolution overlaid were automatically generated by MESSI. Two rankings of substructures based on inferred degrees of coevolution within an alignment of HIV-1 subtype B sequences demonstrated an excess of high ranking substructures that have been commonly cited in the literature as having structure-related importance. This ranking procedure is expected to aid researchers in characterising the secondary structures of less well-studied viruses.

A feature that was not fully accounted for in our model and that is especially important for viral genomes, such as HIV, is that their genomes simultaneously encode for proteins. This implies a dual evolutionary constraint, whereby selection may be acting on the amino acid sequence, whilst simultaneously acting to maintain base-pairing interactions in biologically functional RNA secondary structures. In the future we would like to consider a model that explicitly accounts for both protein-coding and RNA base-pairing constraints.

A second limitation of our model is the assumption of a canonical RNA secondary structure shared across the entire evolutionary history of the sequences being analysed. This is considered a reasonable approximation for low and moderately diverged alignments, where many of the sequences are expected share a high proportion of the same base-pairs. Notwithstanding, it is also likely that different parts of the tree relating the sequences will have at least some parts of those sequence adopting alternative secondary structure conformations. These regions are interesting from a functional perspective. The ability to identify these alternative evolutionary conformations and the mutations responsible for them may lead to significant insights into viral adaptations, such as structural changes following zoonotic transmission of viruses from non-human hosts to humans or the development of drug resistance.

## 5 Software availability

Julia code (compatible with Windows and Linux) is available at: https://github.com/michaelgoldendev/MESSI

## 6 Acknowledgements

MG and OP are supported by the ERC under the European Union’s Seventh Framework Programme (FP7/2007-2013)/ERC grant agreement no. 614725-PATHPHYLODYN.

