## Supplementary material for "Evolutionary analyses of base-pairing interactions in DNA and RNA secondary structures"

### 1 Supplementary methods

#### 1.1 Stationarity and time-reversibility

```
from sympy import *

""" Construct Q matrix """
rates = [Symbol('r1'), Symbol('r2'), Symbol('r3'),
         Symbol('r4'), Symbol('r5'), Symbol('r6')]
freqs = [Symbol('piA'), Symbol('piC'), Symbol('piG'), Symbol('piT')]
Qfreqs = Matrix([freqs[0], freqs[1], freqs[2], freqs[3]])
Q = zeros(4,4)
index = 0
for i in xrange(4):
    for j in xrange(i+1,4):
        if i != j:
            Q[i,j] = freqs[j]*rates[index]
            Q[j,i] = freqs[i]*rates[index]
            index += 1
            if index == 6:
                break
    if index == 6:
            break

""" Calculate Q matrix diagonal entries """
for i in xrange(4):
    Q[i,i]  = 0.0
    for j in xrange(4):
        if i != j:
            Q[i,i] -= Q[i,j]

""" Construct Muse matrix """
Muse = zeros(16,16)
Musefreqs = Matrix([0.0 for i in xrange(16)]) # dinucleotide frequencies

lambdaGC = Symbol('lambdaGC')
lambdaAU = Symbol('lambdaAU')
lambdaGU = Symbol('lambdaGU')
GCindices = [2*4+1,1*4+2] # GC, CG
AUindices = [0*4+3,3*4+0] # AU, UA
GUindices = [2*4+3,3*4+2] # GU, UG

index = 0
kappa = 0 # dinucleotide frequencies normalising constant
for i in xrange(0,4):
    for j in xrange(0,4):
        Musefreqs[index] = Qfreqs[i]*Qfreqs[j]
        dinuc = i*4+j
        if dinuc in GCindices:
            Musefreqs[index] = lambdaGC*lambdaGC*Musefreqs[index]
        if dinuc in AUindices:
```

```python
            Musefreqs[index] = lambdaAU*lambdaAU*Musefreqs[index]
        if dinuc in GUindices:
            Musefreqs[index] = lambdaGU*lambdaGU*Musefreqs[index]
        kappa += Musefreqs[index]
        index += 1

rho1 = Symbol('rho1') # rate for first position
rho2 = Symbol('rho2')  # rate for second position
Musefreqs /= kappa
for i in xrange(16):
    for j in xrange(16):
        if i != j:
            s1 = i / 4
            s2 = i % 4
            e1 = j / 4
            e2 = j % 4
            if s1 != e1 and s2 != e2:
                Muse[i,j] = 0.0
            else:
                if s1 != e1:
                    Muse[i,j] = rho1*Q[s1,e1]
                if s2 != e2:
                    Muse[i,j] = rho2*Q[s2,e2]

            if j in GCindices:
                Muse[i,j] = lambdaGC*Muse[i,j]
            if j in AUindices:
                Muse[i,j] = lambdaAU*Muse[i,j]
            if j in GUindices:
                Muse[i,j] = lambdaGU*Muse[i,j]
            if i in GCindices:
                Muse[i,j] = Muse[i,j]/lambdaGC
            if i in AUindices:
                Muse[i,j] = Muse[i,j]/lambdaAU
            if i in GUindices:
                Muse[i,j] = Muse[i,j]/lambdaGU

""" Calculate Muse matrix diagonal entries """
for i in xrange(0,16):
    Muse[i,i]  = 0.0
    for j in xrange(0,16):
        if i != j:
            Muse[i,i] -= Muse[i,j]

v = simplify(transpose(Musefreqs)*Muse)
print "Stationarity (pi*M = 0)?", (sum([q == 0 for q in v]) == 16)

timereversible = True
for i in xrange(0,16):
    for j in xrange(0,16):
        v = simplify(Musefreqs[i]*Muse[i,j] - Musefreqs[j]*Muse[j,i])
        if v != 0:
            timereversible = False
        #print "pi%d*M%d,%d = pi%d*M%d,%d" % (i,i,j,j,j,i), v == 0
print "Time-reversible (pi_i*M_ij = pi_j*M_ji)?", timereversible

"""
Output:
Stationarity (pi*M = 0)? True
Time-reversible (pi_i*M_ij = pi_j*M_ji)? True
"""
```

Listing 1: Python script (`musesymbolic.py`) testing stationarity and time-reversibility of the extended Muse model.

### 1.2 Modelling site-to-site rate variation

In the M95 and extended M95 models, the substitution rate was assumed to be the same for each of the two nucleotide positions within a pair, as well as across all possible site pairs. However, it is well-established that the rate of substitution can vary across nucleotide positions and that failing to account for rate variability can lead to biased parameter estimates (Yang, 1996). Additionally, many of the datasets analysed in this thesis have coding regions, where it is expected that the third nucleotide position in each codon (the so-called 'codon wobble position') will have relatively higher substitution rates associated with it, due to there being a lower chance of mutations modifying the encoded amino acid.

To model variable substitution rates across sites, the gamma distributed sites rate approach of (Yang, 1993, 1994) was implemented. The GTR model with gamma distributed sites, denoted GTR + Γ, is well-understood for single site substitution models, however, an extension to paired site models had not been previously described. The modified M95 paired site model with gamma distributed site rates, denoted M95 + Γ, is given below:

$$A_{ij} = \begin{cases} \rho_k M_{ij} & \text{if nucleotide pairs } i \text{ and } j \text{ differ in the 1st position,} \\ \rho_l M_{ij} & \text{if nucleotide pairs } i \text{ and } j \text{ differ in the 2nd position,} \\ 0 & \text{2 differences} \end{cases} \quad (1)$$

$\rho_k$ and $\rho_l$ are independently and identically distributed according to a gamma distribution $f(\rho) = \frac{\beta^\alpha \rho^{\alpha-1}}{\Gamma(\alpha)\exp(\beta\rho)}$ with $\mathbb{E}_f[\rho] = \frac{\alpha}{\beta}$. Because $\beta$ is a scale parameter the number of free parameters can be reduced by setting $\alpha = \beta$, which also restricts the mean of the gamma distribution to 1. A small $\alpha$ suggests rates differ significantly across sites, whereas a large $\alpha$ implies roughly equal rates amongst sites.

The conditional probability for a site pair $\mathcal{D}_{q,r}$ with rates $\rho_k$ and $\rho_l$ at sites $\mathcal{D}_q$ and $\mathcal{D}_r$, respectively, is given by $P(\mathcal{D}_{q,r}|\widehat{q,r}, \mathcal{T}, \boldsymbol{\theta}, \rho_k, \rho_l)$ and the marginal probability for a site pair $\mathcal{D}_{q,r}$ is given by:

$$P(\mathcal{D}_{q,r}|\widehat{q,r}, \mathcal{T}, \boldsymbol{\theta}) = \int_0^\infty \int_0^\infty f(\rho_k) f(\rho_l) P(\mathcal{D}_{q,r}|\widehat{q,r}, \mathcal{T}, \boldsymbol{\theta}, \rho_k, \rho_l) dk dl \quad (2)$$

Note that it is necessary to perform a double integration as it is assumed that substitution rates at each of two positions within a nucleotide pair may differ. As is it not computationally feasible to evaluate the double integral in Equation 2 it is approximated using a discrete distribution (Yang, 1993, 1994). Let $F(\rho)$ represent the cumulative distribution function (CDF) of $f(\rho)$ and $F^{-1}(\rho)$ be the inverse CDF, if $p = F(\rho)$ then $\rho = F^{-1}(p)$. The $K$ rate classes are assumed to have equal proportions, i.e. $p_k = \frac{1}{K}$ for all $k$, and the $\rho_k$'s are given by $\rho_k = \frac{F^{-1}(\frac{k}{K+1})}{K^{-1}\sum_{k_2} F^{-1}(\frac{k_2}{K+1})}$ (the denominator ensures that the discretised $\rho_k$'s have a mean of one). Hence, the probability of the data at a particular site pair are approximated by:

$$P(\mathcal{D}_{q,r}|\widehat{q,r}, \lambda, \mathcal{T}, \boldsymbol{\theta}) \approx \sum_{k=1}^{K} \left[ \sum_{l=1}^{K} p_k p_l P(\mathcal{D}_{q,r}|\widehat{q,r}, \lambda, \mathcal{T}, \boldsymbol{\theta}, \rho_k, \rho_l) \right] \quad (3)$$

Note that the double summation in Equation 3 increases the computational time by a factor of $\mathcal{O}(K^2)$ over a paired site model assuming equal substitution rates across sites. For all analyses a discretisation of $K = 3$ was used.

### 1.3 Inside and outside algorithms

Pseudocode for iterative implementations of the inside and outside algorithms are provided in Algorithm S1 and Algorithm S2, respectively.

### 1.4 Sampling secondary structures using the inside probability matrix

Assuming an RNA SCFG written in double emission normal form (Anderson *et al.*, 2012), sampling proceeds recursively starting with the entire alignment (*i.e.* positions 1 to $L$) and the start symbol $S$. The total

**Algorithm S1** Iterative implementation of the inside algorithm for an RNA SCFG in double-emission normal form

```
for U → • in type1rules do
    // consider all type 1 rules (rules generating unpaired positions)

    for q = 1 to L do
        I(U,q,q) = p[U → •] × p(D_q|q̇) // likelihood of column q under the unpaired model
    end
end
for b = 2 to L do
    // consider subsequences of increasing length b

    for r = 1 to L − b + 1 do
        // consider all starting positions r for subsequences of length b

        for U → VW in type3rules do
            // consider all type 3 rules (rules generating a branching)

            for h = r in L + b − 2 do
                // consider all breakpoint positions h generating a branching

                I(U,r,r+b-1) += p[U → VW]
                × I(V,r,h) // probability of the left subsequence being generated by V
                × I(W,h+1,r+b-1) // probability of the right subsequence being generated by W
            end
        end
        for U → (V) in type2rules do
            // consider all type 2 rules (i.e. rules generating a base-pairing between positions r and r + b − 1)

            I(U,r,r+b-1) += p[U → (V)]
            × p(D_{r,r+b−1}|r, r + b − 1 (overarc)) // likelihood of paired sites (r, r + b − 1) under the paired model
            × I(V,r+1,r+b-2) // probability of the subsequence between the base-paired positions being
            generated by V
        end
    end
end
```

**Algorithm S2** Iterative implementation of the outside algorithm for an RNA SCFG in double-emission normal form

```
O(S,1,L) = 1.0
for c = 2 to L do
    b = L − c // consider subsequences of decreasing length b
    for r = 0 to L − b − 1 do
        // consider all starting positions r for subsequences of length b

        for U → VW in type3rules do
            // consider all type 3 rules (rules generating a branching)

            for h = r + b + 1 in L − 1 do
                O(V,r+1,r+b+1) += p[U → VW]
                × O(U,r+1,h+1) // outside probability leading to the generation of U
                × I(W,r+b+2,h+1) // inside probability of the right subsequence
            end
            for h = 0 in r − 1 do
                O(W,r+1,r+b+1) += p[U → VW]
                × O(U,h+1,r+b+1) // outside probability leading to the generation of U
                × I(V,h+1,r) // inside probability of the left subsequence
            end
        end
        for U → (V) in type2rules do
            // consider all type 2 rules (i.e. rules generating a base-pairing between positions r and r + b + 2)

            O(V,r+1,r+b+1) += p[V → (U)]
            × p(D_{r,r+b+2} | r, \widehat{r+b}+2)
            × O(U,r,r+b+2)  // outside probability leading to the generation of U
        end
    end
end
```

probability at this stage is given by element $I(S,1,L)$ of the inside matrix. Each element of the inside matrix, is a sum over event probabilities, each corresponding to one of three possible event types.

(i) a branching event. A position $k$ is sampled along the current sub-alignment leading to the generation of two conditionally independent non-terminal symbols ($L$ and $S$). $L$ and $S$ correspond to left and right sub-alignments, respectively. The left sub-alignment ends at position $k$ and the right sub-alignment starts at position $k+1$. Following the generation of a branching event, the non-terminal symbols $L$ and $S$, corresponding to two new events are recursively sampled.

(ii) an unpairing event. A single terminal symbol, $\bullet$, is emitted at position $k$. As no further non-terminal symbols are generated, further recursive calls are not made.

(iii) a base-pairing event. Terminal symbols '(' and ')' are emitted at the first position, $q$, and last position, $r$, of the current sub-alignment. Furthermore, a non-terminal symbol $F$ corresponding to sub-alignment starting at $q+1$ and ending at $r-1$ between the two base-paired positions is generated and recursively sampled.

Eventually only a string of terminal symbols will remain, at which point the sampled secondary structure can be easily read off.

A code implementation, written in the `Julia` programming language is given in Listing 2. Note that the computational time complexity for sampling a single secondary structures, having precomputed the inside probabilities, is $\mathcal{O}(L^2)$.

```
function samplestructurehelper(rng::AbstractRNG, inside::Array{Float64,3}, pairedlogprobs::Array{Float64,2}, unpairedlogprobs::Array{Float64,1}, parentsymbol::Char, x::Int, y::Int, paired::Array{Int,1}, grammar::KH99, B::Float64)
  type1rules = grammar.type1rules
  type2rules = grammar.type2rules
  type3rules = grammar.type3rules
  stack = Tuple{Char,Int,Int}[]
  push!(stack, (parentsymbol, x, y))
  while length(stack) > 0
    parentsymbol, x, y = pop!(stack)
    rulearr = Rule[]
    if x == y  # unpaired
    elseif x < y
      j = x
      b = y-j+1

      logliks = Float64[]
      rule = Tuple[]

      for type3rule in type3rules  # bifurcations / branching
        if type3rule.left == parentsymbol
          for h=j:j+b-2
            prob1 = inside[type3rule.rightindices[1],j,h]
            prob2 = inside[type3rule.rightindices[2],h+1,j+b-1]
            push!(logliks, type3rule.logprob + prob1 + prob2)
            push!(rule, (3,j,h,h+1,j+b-1,type3rule.right[1],type3rule.right[2]))
            push!(rulearr, type3rule)
          end
        end
      end

      for type2rule in type2rules # base-pairings
        if type2rule.left == parentsymbol
          push!(logliks, (type2rule.logprob + pairedlogprobs[j,j+b-1]) + inside[type2rule.rightindices[2],j+1,j+b-2])
          push!(rule, (2,j,j+b-1,type2rule.right[2]))
          push!(rulearr, type2rule)
        end
      end

      logliks *= B
      s = -Inf
      for v in logliks
        s = logsumexp(s,v)
      end
      r = sample(rng, exp.(logliks-s))  # sample rule used to generate bifurcation or base-pairing

      if rule[r][1] == 2 # base-pairing, recursively sample the sub-alignment
        a = rule[r][2]
        b = rule[r][3]
        paired[a] = b # store pairing
        paired[b] = a # store pairing
        push!(stack, (rule[r][4], a+1, b-1))
      elseif rule[r][1] == 3  # bifurcation, recursively sample the two sub-alignments
        push!(stack, (rule[r][6], rule[r][2], rule[r][3]))
        push!(stack, (rule[r][7], rule[r][4], rule[r][5]))
      end
    end
  end

  return calculateKH99prior(paired)
end
```

Listing 2: Julia recursion (`samplestructurehelper`) for sampling secondary structures from the inside probabilities, phylogenetic unpaired probabilities, and phylogenetic paired probabilities.

### 1.5 Posterior base-pairing and unpairing probabilities

The posterior base-pairing probability of a pair of positions $q$ and $r$ can be obtained by calculating the expected positional emission probabilities of type 2 rules (base-pairing rules) using the inside and outside probabilities:

$$p(\widehat{q,r}|\mathcal{D}) = \frac{\sum_{U \to (V)} O(U,q,r)p[U \to (V)]p(q,r)I(V,q+1,r-1)}{I(S,1,L)} \tag{4}$$

Likewise the unpairing probability of a position $i$ can be obtained using the expected positional emission probabilities of type 1 rules (unpaired rules):

$$p(\dot{q}|\mathcal{D}) = \frac{\sum_{U \to \bullet} O(U,q,q)p[U \to \bullet]p(q)}{I(S,1,L)} \tag{5}$$

### 1.6 Consensus structure prediction

The maximum expected accuracy (MEA) algorithm was used to find the secondary structure with the highest sum of base-pairing and unpaired probabilities (Lu *et al.*, 2009). The MEA value and secondary structure is obtained by computing the dynamic programming matrix:

$$E_{q,r} = \max \begin{cases} p(\dot{q}|\mathcal{D}) + E_{q+1,r} \\ \alpha p(\widehat{q,r}|\mathcal{D}) + E_{q+1,r-1} \\ \max\limits_{q+1<k<r} \alpha p(\widehat{q,k}|\mathcal{D}) + E_{q+1.k-1} + E_{k+1,r} \end{cases} \tag{6}$$

Where $E_{q,r} := 0$ for $q \geq r$, the MEA value is given by $E_{1,L}$, and the MEA structure is obtained via backtracking through the matrix, starting with $E_{1,L}$. The parameter, $\alpha = 2$, controls the trade-off between recall and precision in predicting base-pairs, where higher $\alpha$'s encourage a greater number of base-pairings to be predicted over unpaired positions.

### 1.7 Datasets

Three classes of datasets were analysed. The first class of datasets consisted of non-coding RNA alignments obtained from the RFAM database (Burge *et al.*, 2012). Each RFAM dataset has an associated consensus secondary structure based on experimental RNA structure determination. These consensus secondary structures were only used for secondary structure prediction benchmarking purposes and were not conditioned on during inference. RFAM datasets are denoted with 'RF' prefix in their name. The second and third classes of datasets analysed, consisted of the complete genomes of single-stranded RNA and single-stranded DNA viruses, respectively, obtained from the NCBI nucleotide database (Acland *et al.*, 2014) and aligned using MUSCLE (Edgar, 2004). Typically these datasets did not have associated consensus secondary structures.

#### 1.7.1 Representative subset selection

Many of the viruses selected for analysis had a large number of complete genomic sequences available ($200-2000$ sequences) in the NCBI nucleotide database. Because the inference algorithms scale linearly in the number of sequences, we chose to reduce each full dataset $F$ to a smaller representative subset $S^*$ of $N$ sequences, where $N$ is typically 100-200 sequences. A genetic algorithm was used to find the optimal (or near-optimal) subset $S^*$ containing $N$ sequences such that following criteria was maximised:

$$S^* = \operatorname*{argmax}_{S \subseteq F, |S|=N} \sigma(S) \tag{7}$$

where

$$\sigma(S) = \sum_{s \in S} \min_{s' \in S \setminus s} d(s, s') \tag{8}$$

where $d(s, s')$ is the hamming distance between aligned sequences $s$ and $s'$.

In other words: $S^*$ was chosen amongst possible subsets in $F$ containing $N$ sequences such that the sum of the distances between each sequence and its closest sequence in the subset was maximised. The result is a subset $S^*$ that attempts to maximise the amount of diversity (*i.e.* information) in each dataset, whilst reducing the number of sequences used during inference, to help reduce the computational burden.

##### 1.7.2 Linearisation of circular DNA viruses

Most known DNA viruses have circular genomes - genomes that form a closed loop without 5' and 3' ends. This presents a problem when modelling their secondary structures, as the underlying SCFG and inside-outside algorithm assume linear sequences.

To address this, we linearised every circular DNA virus alignment by choosing an appropriate point of linearisation. We chose to linearise the DNA virus alignments at their aligned origin of replication (*ori*) sites. All of the DNA viruses in our analysis contain known ori's that form highly stable secondary structures (Muhire *et al.*, 2014). We linearise each alignment within the loop of region of these structures, thereby ensuring that long-range base-pairings are maintained.

#### 1.8 Measures of predictive accuracy

To measure the predictive accuracy of a secondary structure prediction it is necessary to have a *reference* secondary structure (typically determined from experiment) to measure the prediction against. In Figure S1 we give an example of a predicted structure and a reference structure and use them to compare different measures of predictive accuracy in this section.

The first few measures described here require counts of *true positives* (TP, the number of correctly predicted base-pairs), *false negatives* (FN, the number of base-pairs in the reference structure not in the predicted structure), and *false positives* (FP, the number of base-pairs predicted not in the reference structure). The number of *true negatives* (TN) is also required in some cases, which is defined as $\text{TN} = \frac{L(L-1)}{2} - \text{TP} - \text{FN} - \text{FP}$. Finally, the number of compatible base-pairs, defined as the number of FP base-pairs compatible with all base-pairs in the reference structure, is also sometimes used to reduce the effect false positives when measuring predictive accuracy.

##### 1.8.1 Precision, recall, and F-score

Two simple measures of predictive accuracy are precision and recall (Gardner and Giegerich, 2004), defined as follows:

$$\text{Precision} = \frac{\text{TP}}{\text{TP} + \text{FP}} \tag{9}$$

$$\text{Recall} = \frac{\text{TP}}{\text{TP} + \text{FN}} \tag{10}$$

Precision can be interpret as the number of correctly predicted base-pairings (TP) compared to the total number of predicted base-pairings (TP + FP), whereas recall can be interpreted as the number of correctly predicted base-pairings (TP) compared to the total number of base-pairings in the reference structure (TP + FN).

Note that recall can be increased by encouraging a particular prediction method to predict more base-pairings; however, this will likely result in an increase in the number false positives (FP) and a corresponding decrease in precision. Therefore any particular method should aim to score high on both metrics, for this reason the two metrics are often "averaged" by taking their harmonic mean (termed the F1-score) as follows:

$$\text{F1 score} = 2 \cdot \frac{\text{precision} \cdot \text{recall}}{\text{precision} + \text{recall}} \tag{11}$$

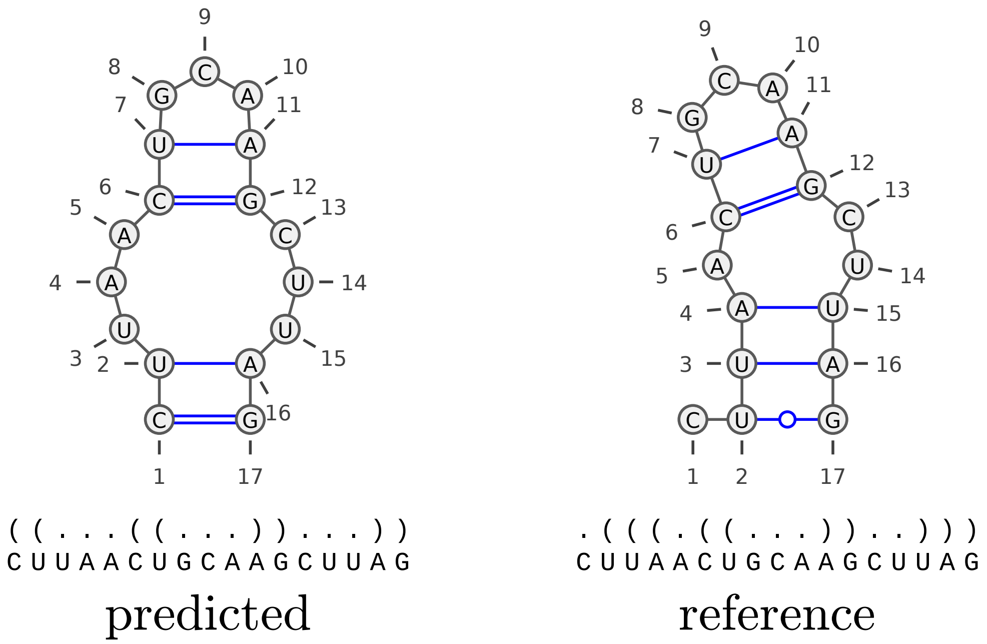


Figure S1: Two example secondary structures visualised using VARNA (Darty *et al.*, 2009) with the corresponding dot bracket structures indicated below. The first is a 'predicted' structure and the second is a 'reference' structure corresponding to the same sequence. True positive base-pairs (TP=2), false negative base-pairs (FN=3), false positive base-pairs (FP=2), true negative base-pairs (TN $= \frac{L(L-1)}{2} - \text{TP} - \text{FN} - \text{FP} = 136 - 2 - 3 - 2 = 129$), and compatible bas-pairs ($\xi = 2$).

#### 1.8.2 Matthews correlation coefficient

Another commonly used measure of predictive accuracy is Matthews correlation coefficient (MCC; Baldi *et al.* (2000)). Like the F-score, MCC combines both precision and recall in a single measure. It is defined as follows:

$$\text{MCC} = \frac{\text{TP} \times \text{TN} - (\text{FP} - \xi) \times \text{FN}}{\sqrt{(\text{TP} + \text{FP} - \xi)(\text{TP} + \text{FN})(\text{TN} + \text{FP} - \xi)(\text{TN} + \text{FN})}} \tag{12}$$

MCC ranges in value from -1 (completely wrong) to 1 (perfect prediction). However, in the case of secondary structure, MCC typically ranges from 0 to 1, because TN= 0 rarely occurs in practice (Gardner and Giegerich, 2004).

#### 1.8.3 Mountain metric

The mountain metric is a measure of secondary structure distance based on the mountain representation of secondary structures, which follows naturally from the dot-bracket representation of secondary structures.

Using the mountain representations for secondary structures it is simple to graphically derive the mountain metric distance (see Figure S2 for an explanation). For a complete mathematical exposition see Moulton *et al.* (2000).

The mountain distance provides a nuanced measure of structural similarity compared to the simple base-pair metrics described above. For example, it better captures structural features compared to base-pair metrics that heavily penalize base-pairings that are even slightly offset relative to one another. For example: shifting the base-pairings, as in Figure S2, would be heavily penalized by simple base-pair metrics (resulting in an F1 score of 0.44 or 44% in this example), whereas the mountain similarity metric better captures similarities in shape (resulting in a mountain similarity measure of 0.819 or 81.9%).

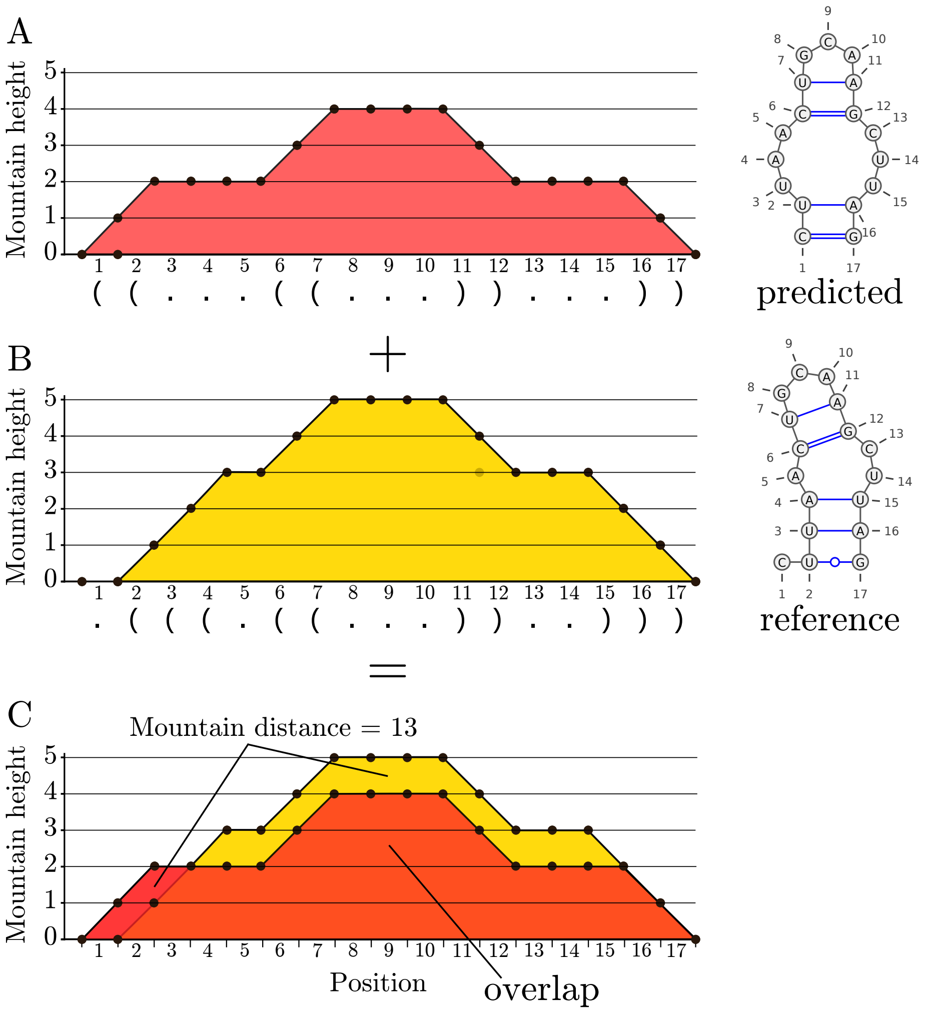


Mountain diameter = 72

Normalised mountain distance = 13 / 72 = 0.181

Mountain similarity = 1 - 0.181 = 0.819 or 81.9%

Figure S2: Mountain plots of two secondary structures. The mountain height at a particular sequence position corresponds to the number of base-pairs that enclose that position. For example, immediately after the third position in both structures (plots A and B) the mountain height is 2, because both positions are enclosed by two base-pairs. The non-overlapping areas in red and yellow (plot C) provide a graphical depiction of the mountain distance between the two secondary structures. The mountain distance can be normalised by the mountain diameter. The mountain diameter is the maximum possible mountain distance between two structures of a given length.

The mountain metric together with a set of secondary structures defines a *metric space*. A metric space consists of a set and a distance function, where the distance function has certain properties. One property is symmetry, that is $d(x, y) = d(y, x)$ - the distance remains the same regardless of the ordering of the two elements being compared. Note that symmetry does not hold for measures such as precision, recall, or MCC, and hence these measures can not be used as distance functions as part of metric spaces.

#### 1.9 Substructure ranking

To rank substructures, the complete secondary structure of a nucleic acid sequence was subdivided into a number of smaller substructures 10-350 nucleotides in length. The substructures were then ranked by the numeric values within them. The ranking was performed by comparing two lists of numeric values: one corresponding to values within the substructure, and one corresponding to all values outside of the substructure (all values in the complete structure, but not in the substructure).

The two lists were compared using a Wilcoxon-rank sum test which assesses statistically whether values within the substructure are significantly higher (or lower) than values outside of it. The Wilcoxon-rank test statistic is used to compute a z-score for each substructure indicating the degree of difference. For example, a substructure with a large positive z-score would suggest that the distribution of numeric values within the substructure are significantly higher than the distribution of numeric values outside of the substructure. By using a Wilcoxon-rank sum test, this method attempts to account for the number of data points associated with each substructure. Substructures are subsequently ranked by sorting by their z-scores.

### 2 Supplementary results

Table S1: Maximum log-likelihood (MLL), structure information entropy ($H$), maximum information entropy ($H_{\max}$), and normalised structure information entropy ($H/H_{\max}$) values for nine different datasets and for two random permutations ($p_1$ and $p_2$) of those datasets.

| Dataset | MLL | ΔMLL unpaired | Length | $H$ | $H_{\max}$ | $H/H_{\max}$ | ΔMLL $p_1$ | $\Delta H$ $p_1$ | $\Delta H/H_{\max}$ $p_1$ | ΔMLL $p_2$ | $\Delta H$ $p_2$ | $\Delta H/H_{\max}$ $p_2$ |
|---|---|---|---|---|---|---|---|---|---|---|---|---|
| WDV | -10731.6 | -88.8 | 2755 | 1187.7 | 3806.9 | 0.312 | -50.8 | +276.6 | +0.073 | -56.4 | +365.3 | +0.096 |
| MSV | -15341.5 | -270.6 | 2755 | 1484.9 | 3806.9 | 0.390 | -245.9 | +99.3 | +0.026 | -234.8 | +10.7 | +0.003 |
| RF00001 | -27897.7 | -3605.0 | 230 | 1.4 | 307.6 | 0.005 | -2509.6 | +6.4 | +0.021 | -2539.9 | +6.2 | +0.020 |
| RF00002 | -4399.1 | -162.5 | 207 | 29.2 | 275.9 | 0.106 | -115.4 | +2.7 | +0.010 | -100.6 | +13.1 | +0.047 |
| RF00003 | -4653.0 | -322.5 | 203 | 14.7 | 270.4 | 0.054 | -255.8 | -0.3 | -0.001 | -268.5 | -3.6 | -0.013 |
| RF00004 | -16716.6 | -1733.6 | 278 | 7.9 | 373.8 | 0.021 | -1085.6 | +7.6 | +0.020 | -1301.3 | +18.0 | +0.048 |
| Hepatitis A | -63755.3 | -434.3 | 7572 | 3327.1 | 10490.7 | 0.317 | -221.0 | +720.0 | +0.069 | -200.1 | +586.8 | +0.056 |
| Rhinovirus A | -217136.2 | -6905.2 | 7308 | 1524.4 | 10124.4 | 0.151 | -1118.7 | +135.1 | +0.013 | -1120.7 | +127.6 | +0.013 |
| HPV1 | -64966.3 | -438.5 | 7668 | 3023.8 | 10624.0 | 0.285 | -281.6 | +541.7 | +0.051 | -306.4 | +810.8 | +0.076 |

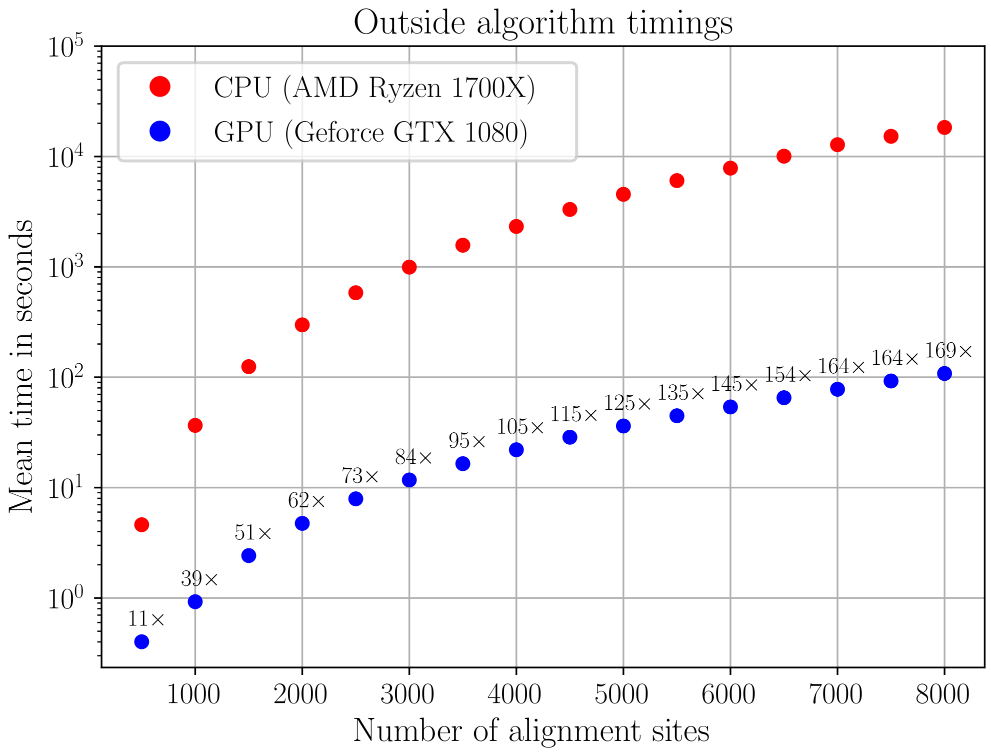


Figure S3: Outside algorithm timings in seconds ($\log_{10}$ axis) as a function of the number of alignment sites, with the fold speed-up of the GPU version over the CPU version indicated above the GPU timings.

Table S2: Secondary structure prediction benchmarks of 22 RFAM datasets.

| Dataset | Method | Precision | Recall | F-score | MCC | Weighted mountain similarity | Mountain similarity (p=1) |
|---|---|---|---|---|---|---|---|
| RF01854 | Our method | 0.871 | **0.670** | **0.758** | **0.781** | **0.856** | **0.936** |
| | PPfold | 0.833 | 0.275 | 0.413 | 0.524 | 0.611 | 0.687 |
| | RNAalifold | **1.000** | 0.319 | 0.483 | 0.564 | 0.605 | 0.692 |
| RF01846 | Our method | 0.856 | **0.975** | **0.911** | **0.987** | **0.958** | 0.961 |
| | PPfold | 0.889 | 0.608 | 0.722 | 0.779 | 0.907 | **0.978** |
| | RNAalifold | **0.943** | 0.418 | 0.579 | 0.646 | 0.857 | 0.967 |
| RF00002 | Our method | 0.792 | **0.760** | **0.776** | **0.872** | **0.907** | **0.991** |
| | PPfold | 0.679 | 0.760 | 0.717 | 0.872 | 0.898 | 0.980 |
| | RNAalifold | **1.000** | 0.480 | 0.649 | 0.693 | 0.874 | 0.952 |
| RF02001 | Our method | 0.872 | **0.854** | 0.863 | **0.924** | **0.966** | **0.963** |
| | PPfold | 0.930 | 0.833 | **0.879** | 0.913 | 0.962 | 0.947 |
| | RNAalifold | **0.968** | 0.625 | 0.759 | 0.790 | 0.907 | 0.914 |
| RF00379 | Our method | 0.424 | **0.875** | 0.571 | 0.825 | 0.898 | 0.868 |
| | PPfold | **0.519** | 0.875 | **0.651** | **0.825** | **0.934** | 0.893 |
| | RNAalifold | 0.519 | 0.875 | 0.651 | 0.825 | 0.934 | **0.904** |
| RF00380 | Our method | 0.469 | 0.920 | 0.622 | 0.693 | 0.826 | 0.742 |
| | PPfold | 0.533 | **0.960** | 0.686 | **0.732** | 0.855 | 0.774 |
| | RNAalifold | **0.548** | 0.920 | **0.687** | 0.718 | **0.877** | **0.789** |
| RF00012 | Our method | 0.886 | **0.984** | **0.932** | **0.992** | **0.955** | **0.982** |
| | PPfold | 0.784 | 0.635 | 0.702 | 0.797 | 0.890 | 0.977 |
| | RNAalifold | **1.000** | 0.603 | 0.752 | 0.776 | 0.841 | 0.932 |
| RF00100 | Our method | 0.902 | 0.632 | 0.743 | 0.795 | 0.854 | 0.964 |
| | PPfold | 0.903 | **0.644** | **0.752** | **0.802** | **0.862** | **0.969** |
| | RNAalifold | **0.921** | 0.402 | 0.560 | 0.634 | 0.744 | 0.933 |
| RF00020 | Our method | 0.811 | **1.000** | 0.896 | **1.000** | 0.926 | 0.957 |
| | PPfold | 0.879 | 0.967 | **0.921** | 0.983 | **0.947** | **0.988** |
| | RNAalifold | **1.000** | 0.700 | 0.824 | 0.836 | 0.905 | 0.980 |
| RF00174 | Our method | 0.544 | 0.861 | **0.667** | **0.817** | 0.903 | 0.901 |
| | PPfold | 0.507 | **0.972** | 0.667 | 0.786 | 0.850 | 0.801 |
| | RNAalifold | **0.750** | 0.583 | 0.656 | 0.661 | **0.945** | **0.960** |

| Dataset | Method | Precision | Recall | F-score | MCC | Weighted mountain similarity | Mountain similarity (p=1) |
|---|---|---|---|---|---|---|---|
| RF00001 | Our method | 0.707 | **0.853** | **0.773** | **0.923** | 0.933 | 0.973 |
| | PPfold | 0.714 | 0.735 | 0.725 | 0.857 | **0.964** | **0.991** |
| | RNAalifold | **0.840** | 0.618 | 0.712 | 0.786 | 0.910 | 0.971 |
| RF00004 | Our method | 0.830 | **0.978** | 0.898 | **0.989** | 0.928 | 0.982 |
| | PPfold | 0.913 | 0.933 | 0.923 | 0.966 | 0.949 | **0.996** |
| | RNAalifold | **1.000** | 0.889 | **0.941** | 0.943 | **0.964** | 0.994 |
| RF00030 | Our method | 0.653 | **0.721** | **0.685** | **0.849** | **0.984** | 0.990 |
| | PPfold | 0.642 | 0.632 | 0.637 | 0.795 | 0.978 | **0.991** |
| | RNAalifold | **0.706** | 0.529 | 0.605 | 0.728 | 0.970 | 0.989 |
| RF02542 | Our method | 0.641 | **0.863** | **0.735** | **0.929** | 0.863 | 0.957 |
| | PPfold | 0.637 | 0.758 | 0.692 | 0.871 | 0.875 | 0.964 |
| | RNAalifold | **0.698** | 0.698 | 0.698 | 0.836 | **0.889** | **0.970** |
| RF00026 | Our method | 0.250 | **1.000** | 0.400 | 0.913 | 0.860 | 0.976 |
| | PPfold | 0.385 | 1.000 | 0.556 | **1.000** | 0.925 | 0.987 |
| | RNAalifold | **1.000** | 1.000 | **1.000** | 1.000 | **1.000** | **1.000** |
| RF00017 | Our method | 0.962 | **0.850** | **0.903** | **0.922** | **0.902** | **0.938** |
| | PPfold | 0.959 | 0.783 | 0.862 | 0.885 | 0.883 | 0.908 |
| | RNAalifold | **0.987** | 0.650 | 0.784 | 0.806 | 0.800 | 0.861 |
| RF00011 | Our method | 0.664 | **0.987** | 0.794 | **0.974** | 0.877 | 0.909 |
| | PPfold | 0.634 | 0.934 | 0.755 | 0.947 | 0.877 | 0.918 |
| | RNAalifold | **0.767** | 0.908 | **0.831** | 0.953 | **0.948** | **0.945** |
| RF00010 | Our method | 0.617 | **0.987** | **0.759** | **0.891** | 0.909 | 0.912 |
| | PPfold | 0.591 | 0.907 | 0.716 | 0.847 | 0.918 | 0.912 |
| | RNAalifold | **0.667** | 0.720 | 0.692 | 0.740 | **0.960** | **0.935** |
| RF00209 | Our method | 0.857 | 0.934 | 0.894 | 0.967 | **0.950** | **0.993** |
| | PPfold | 0.844 | **0.975** | **0.905** | **0.988** | 0.933 | 0.981 |
| | RNAalifold | **0.877** | 0.877 | 0.877 | 0.936 | 0.939 | 0.988 |
| RF02540 | Our method | 0.650 | **0.933** | 0.766 | **0.958** | 0.858 | 0.971 |
| | PPfold | 0.677 | 0.908 | 0.775 | 0.946 | 0.882 | 0.973 |
| | RNAalifold | **0.768** | 0.890 | **0.825** | 0.940 | **0.927** | **0.982** |

| Dataset | Method | Precision | Recall | F-score | MCC | Weighted mountain similarity | Mountain similarity (p=1) |
|---|---|---|---|---|---|---|---|
| RF02541 | Our method | 0.677 | **0.897** | 0.772 | **0.941** | 0.902 | 0.978 |
| | PPfold | 0.707 | 0.884 | 0.786 | 0.934 | 0.915 | 0.980 |
| | RNAalifold | **0.803** | 0.852 | **0.827** | 0.917 | **0.939** | **0.986** |
| RF00003 | Our method | 0.800 | **0.900** | 0.847 | 0.949 | 0.911 | 0.976 |
| | PPfold | **0.900** | 0.900 | **0.900** | **0.949** | **0.948** | **0.986** |
| | RNAalifold | 0.897 | 0.875 | 0.886 | 0.935 | 0.931 | 0.982 |

Table S3: SHAPE structure ranking. 86 non-overlapping HIV NL4-3 substructures ranked from highest to lowest z-score based on the estimated degrees of coevolution within an alignment of HIV-1 subtype B sequences. Where the HIV NL4-3 SHAPE secondary structure was used as the canonical structure.

| Rank | Alignment position | Mapped position | Length | Name and reference | Median | z-score |
|---|---|---|---|---|---|---|
| 1 | 8233 - 8582 | 7249 - 7595 | 350 | Rev Response element (RRE, Heaphy *et al.* (1990); Mandal and Breaker (2004)) | 5.38 | 5.02 |
| 2 | 2608 - 2943 | 1991 - 2326 | 336 | Longest continuous helix (Siegfried *et al.*, 2014) | 5.17 | 2.92 |
| 3 | 10155 - 10383 | 8982 - 9170 | 229 | 3’ Untranslated region (3’UTR, Siegfried *et al.* (2014)) | 5.27 | 2.69 |
| 4 | 588 - 838 | 105 - 344 | 251 | 5’ Untranslated region (5’UTR, Siegfried *et al.* (2014)) | 5.65 | 2.61 |
| 5 | 9570 - 9584 | 8440 - 8454 | 15 | | 5.91 | 2.29 |
| 6 | 860 - 979 | 366 - 485 | 120 | 5’ Untranslated region (5’UTR, Siegfried *et al.* (2014)) | 5.54 | 2.28 |
| 7 | 1710 - 1845 | 1177 - 1312 | 136 | | 5.17 | 2.28 |
| 8 | 2115 - 2301 | 1561 - 1711 | 187 | Gag-pol frameshift (Chamorro *et al.*, 1992) | 5.31 | 2.21 |
| 9 | 1479 - 1490 | 946 - 957 | 12 | | 5.85 | 2.04 |
| 10 | 3886 - 3907 | 3269 - 3290 | 22 | | 5.80 | 2.01 |
| 11 | 1404 - 1446 | 871 - 913 | 43 | | 5.77 | 1.95 |
| 12 | 1559 - 1601 | 1026 - 1068 | 43 | | 5.65 | 1.84 |
| 13 | 4716 - 4746 | 4099 - 4129 | 31 | | 5.68 | 1.80 |
| 14 | 550 - 587 | 67 - 104 | 38 | 5’ Untranslated region (5’UTR, Siegfried *et al.* (2014)) | 5.72 | 1.62 |
| 15 | 1454 - 1463 | 921 - 930 | 10 | | 5.80 | 1.59 |
| 16 | 995 - 1247 | 501 - 714 | 253 | 5’ Untranslated region (5’UTR, Siegfried *et al.* (2014)) | 5.17 | 1.58 |
| 17 | 4793 - 5090 | 4176 - 4473 | 298 | Central polypurine tract (CPPT, Siegfried *et al.* (2014)) | 5.17 | 1.54 |
| 18 | 1494 - 1547 | 961 - 1014 | 54 | | 5.46 | 1.44 |
| 19 | 3093 - 3149 | 2476 - 2532 | 57 | | 5.40 | 1.40 |
| 20 | 9630 - 9920 | 8500 - 8783 | 291 | | 5.17 | 1.35 |
| 21 | 1908 - 1920 | 1375 - 1387 | 13 | | 5.17 | 1.15 |
| 22 | 4775 - 4792 | 4158 - 4175 | 18 | | 5.80 | 1.04 |
| 23 | 1328 - 1382 | 795 - 849 | 55 | | 5.22 | 0.94 |
| 24 | 5627 - 5639 | 5010 - 5022 | 13 | | 5.45 | 0.93 |
| 25 | 4414 - 4675 | 3797 - 4058 | 262 | | 5.17 | 0.90 |
| 26 | 4685 - 4715 | 4068 - 4098 | 31 | | 5.17 | 0.81 |
| 27 | 485 - 539 | 2 - 56 | 55 | 5’ Trans-activation response element (5’ TAR, Roy *et al.* (1990)) | 5.34 | 0.74 |
| 28 | 3657 - 3675 | 3040 - 3058 | 19 | | 5.77 | 0.72 |
| 29 | 6712 - 6733 | 6048 - 6066 | 22 | | 5.37 | 0.70 |
| 30 | 1608 - 1634 | 1075 - 1101 | 27 | | 5.17 | 0.68 |
| 31 | 2427 - 2535 | 1810 - 1918 | 109 | | 5.34 | 0.67 |
| 32 | 10144 - 10153 | 8971 - 8980 | 10 | | 5.80 | 0.66 |
| 33 | 3795 - 3807 | 3178 - 3190 | 13 | | 5.17 | 0.64 |
| 34 | 1849 - 1881 | 1316 - 1348 | 33 | | 5.17 | 0.63 |
| 35 | 3859 - 3874 | 3242 - 3257 | 16 | | 5.17 | 0.50 |
| 36 | 7391 - 7413 | 6518 - 6540 | 23 | | 5.17 | 0.47 |
| 37 | 5555 - 5616 | 4938 - 4999 | 62 | | 5.17 | 0.38 |

| Rank | Alignment position | Mapped position | Length | Name and reference | Median | z-score |
|---|---|---|---|---|---|---|
| 38 | 5137 - 5146 | 4520 - 4529 | 10 | | 4.96 | 0.38 |
| 39 | 9177 - 9213 | 8169 - 8205 | 37 | | 4.97 | 0.27 |
| 40 | 5726 - 6073 | 5106 - 5445 | 348 | | 5.15 | 0.17 |
| 41 | 5092 - 5108 | 4475 - 4491 | 17 | | 5.17 | 0.12 |
| 42 | 5190 - 5203 | 4573 - 4586 | 14 | | 4.54 | 0.07 |
| 43 | 3154 - 3203 | 2537 - 2586 | 50 | | 5.17 | 0.06 |
| 44 | 7543 - 7557 | 6667 - 6678 | 15 | | 2.98 | 0.06 |
| 45 | 3691 - 3706 | 3074 - 3089 | 16 | | 3.03 | 0.01 |
| 46 | 7363 - 7374 | 6490 - 6501 | 12 | | 4.44 | -0.01 |
| 47 | 2995 - 3046 | 2378 - 2429 | 52 | | 4.86 | -0.03 |
| 48 | 3589 - 3655 | 2972 - 3038 | 67 | | 4.78 | -0.10 |
| 49 | 5205 - 5551 | 4588 - 4934 | 347 | | 5.05 | -0.14 |
| 50 | 2318 - 2372 | 1728 - 1758 | 55 | | 3.87 | -0.31 |
| 51 | 6745 - 6763 | 6078 - 6096 | 19 | | 4.20 | -0.31 |
| 52 | 3924 - 3968 | 3307 - 3351 | 45 | | 4.25 | -0.46 |
| 53 | 1636 - 1674 | 1103 - 1141 | 39 | | 4.60 | -0.50 |
| 54 | 3709 - 3793 | 3092 - 3176 | 85 | | 4.35 | -0.63 |
| 55 | 1292 - 1308 | 759 - 775 | 17 | | 4.73 | -0.65 |
| 56 | 7056 - 7070 | 6273 - 6287 | 15 | | 0.06 | -0.72 |
| 57 | 9234 - 9276 | 8226 - 8268 | 43 | | 4.86 | -0.72 |
| 58 | 1928 - 2113 | 1395 - 1559 | 186 | | 4.56 | -0.83 |
| 59 | 7492 - 7526 | 6616 - 6650 | 35 | | 2.98 | -0.85 |
| 60 | 3822 - 3840 | 3205 - 3223 | 19 | | 3.86 | -0.86 |
| 61 | 8589 - 8612 | 7602 - 7613 | 24 | | 2.95 | -0.86 |
| 62 | 7417 - 7465 | 6544 - 6589 | 49 | | 3.03 | -1.00 |
| 63 | 9591 - 9628 | 8461 - 8498 | 38 | | 3.50 | -1.21 |
| 64 | 3984 - 4327 | 3367 - 3710 | 344 | | 5.07 | -1.27 |
| 65 | 6448 - 6661 | 5793 - 6000 | 214 | SP stem (Siegfried *et al.* (2014)) | 4.65 | -1.34 |
| 66 | 6101 - 6446 | 5473 - 5791 | 346 | | 4.58 | -1.43 |
| 67 | 3219 - 3544 | 2602 - 2927 | 326 | | 4.95 | -1.43 |
| 68 | 7659 - 7684 | 6777 - 6802 | 26 | | 0.38 | -1.43 |
| 69 | 5701 - 5716 | 5081 - 5096 | 16 | | 2.37 | -1.51 |
| 70 | 2588 - 2603 | 1971 - 1986 | 16 | | 2.30 | -1.60 |
| 71 | 9283 - 9391 | 8275 - 8348 | 109 | | 2.60 | -1.62 |
| 72 | 8613 - 8650 | 7614 - 7645 | 38 | | 3.46 | -1.70 |
| 73 | 7559 - 7581 | 6680 - 6702 | 23 | | 1.60 | -1.77 |
| 74 | 7585 - 7649 | 6706 - 6767 | 65 | | 0.22 | -1.90 |
| 75 | 8652 - 8697 | 7647 - 7692 | 46 | | 2.69 | -1.91 |
| 76 | 2374 - 2399 | 1760 - 1785 | 26 | | 2.70 | -2.01 |
| 77 | 8710 - 8775 | 7705 - 7770 | 66 | | 0.23 | -2.22 |
| 78 | 8830 - 9169 | 7825 - 8161 | 340 | | 4.62 | -2.24 |

| Rank | Alignment position | Mapped position | Length | Name and reference | Median | z-score |
|---|---|---|---|---|---|---|
| 79 | 9934 - 10054 | 8797 - 8917 | 121 | | 2.77 | -2.32 |
| 80 | 7789 - 8044 | 6892 - 7075 | 256 | | 3.66 | -2.36 |
| 81 | 7085 - 7361 | 6302 - 6488 | 277 | | 3.63 | -2.46 |
| 82 | 8046 - 8080 | 7077 - 7111 | 35 | | 0.49 | -2.81 |
| 83 | 7766 - 7784 | 6869 - 6887 | 19 | | 0.06 | -2.93 |
| 84 | 8082 - 8106 | 7113 - 7137 | 25 | | 1.16 | -3.16 |
| 85 | 9397 - 9420 | 8354 - 8377 | 24 | | 0.01 | -3.37 |
| 86 | 6800 - 7055 | 6133 - 6272 | 256 | | 0.15 | -6.54 |

Table S4: Consensus structure ranking. 118 non-overlapping HIV consensus substructures ranked from highest to lowest z-score based on their degrees of coevolution within an alignment of HIV-1 subtype B sequences. Where the canonical structure was treated as unknown and a consensus structure predicted

| Rank | Alignment position | Mapped position | Length | Name and reference | Median | z-score |
|---|---|---|---|---|---|---|
| 1 | 8240 - 8577 | 7256 - 7590 | 338 | Rev Response element (RRE, Heaphy *et al.* (1990); Mandal and Breaker (2004)) | 5.64 | 6.53 |
| 2 | 2202 - 2229 | 1645 - 1672 | 28 | Gag-pol frameshift (Chamorro *et al.*, 1992) | 8.17 | 4.56 |
| 3 | 1710 - 1845 | 1177 - 1312 | 136 | | 6.44 | 4.50 |
| 4 | 4751 - 4833 | 4134 - 4216 | 83 | | 6.47 | 3.97 |
| 5 | 4505 - 4709 | 3888 - 4092 | 205 | | 5.22 | 3.21 |
| 6 | 591 - 939 | 108 - 445 | 349 | 5' Untranslated region (5'UTR, Siegfried *et al.* (2014)) | 5.38 | 3.16 |
| 7 | 133 - 151 | NA | 19 | | 6.85 | 2.94 |
| 8 | 2564 - 2890 | 1947 - 2273 | 327 | Longest continuous helix (Siegfried *et al.*, 2014) | 4.44 | 2.62 |
| 9 | 9782 - 9800 | 8645 - 8663 | 19 | | 6.92 | 2.55 |
| 10 | 3612 - 3623 | 2995 - 3006 | 12 | | 6.74 | 2.50 |
| 11 | 5690 - 5720 | 5070 - 5100 | 31 | | 5.76 | 2.43 |
| 12 | 9873 - 9881 | 8736 - 8744 | 9 | | 10.71 | 2.31 |
| 13 | 5622 - 5642 | 5005 - 5025 | 21 | | 7.22 | 2.26 |
| 14 | 9733 - 9781 | 8597 - 8644 | 49 | | 5.20 | 2.22 |
| 15 | 10008 - 10020 | 8871 - 8883 | 13 | | 7.39 | 2.00 |
| 16 | 10061 - 10390 | NA | 330 | 3' Untranslated region (3'UTR, Siegfried *et al.* (2014)) | 4.54 | 1.82 |
| 17 | 7410 - 7425 | 6537 - 6552 | 16 | | 5.65 | 1.78 |
| 18 | 1185 - 1211 | NA | 27 | | 5.63 | 1.76 |
| 19 | 3538 - 3552 | 2921 - 2935 | 15 | | 5.97 | 1.50 |
| 20 | 8057 - 8214 | 7088 - 7230 | 158 | | 4.77 | 1.49 |
| 21 | 1080 - 1146 | 586 - 652 | 67 | 5' Untranslated region (5'UTR, Siegfried *et al.* (2014)) | 6.49 | 1.45 |
| 22 | 6627 - 6643 | 5966 - 5982 | 17 | SP stem (Siegfried *et al.* (2014)) | 4.98 | 1.44 |
| 23 | 3261 - 3268 | 2644 - 2651 | 8 | | 5.79 | 1.28 |
| 24 | 3577 - 3586 | 2960 - 2969 | 10 | | 5.64 | 1.09 |
| 25 | 7434 - 7468 | NA | 35 | | 4.99 | 1.00 |
| 26 | 105 - 118 | NA | 14 | | 4.82 | 0.86 |
| 27 | 9443 - 9464 | NA | 22 | | 5.35 | 0.80 |
| 28 | 7536 - 7648 | 6660 - 6766 | 113 | | 4.63 | 0.77 |
| 29 | 1866 - 1882 | 1333 - 1349 | 17 | | 4.98 | 0.77 |
| 30 | 4161 - 4187 | 3544 - 3570 | 27 | | 4.41 | 0.66 |
| 31 | 8005 - 8054 | 7036 - 7085 | 50 | | 4.39 | 0.52 |
| 32 | 3342 - 3393 | 2725 - 2776 | 52 | | 3.98 | 0.52 |
| 33 | 3270 - 3299 | 2653 - 2682 | 30 | | 4.93 | 0.48 |
| 34 | 1908 - 1920 | 1375 - 1387 | 13 | | 4.28 | 0.28 |
| 35 | 4306 - 4320 | 3689 - 3703 | 15 | | 4.02 | 0.26 |
| 36 | 953 - 961 | 459 - 467 | 9 | 5' Untranslated region (5'UTR, Siegfried *et al.* (2014)) | 4.56 | 0.19 |
| 37 | 1422 - 1707 | 889 - 1174 | 286 | | 4.23 | 0.14 |

| Rank | Alignment position | Mapped position | Length | Name and reference | Median | z-score |
|---|---|---|---|---|---|---|
| 38 | 419 - 476 | NA | 58 | | 3.98 | 0.11 |
| 39 | 7918 - 7923 | 6982 - 6987 | 6 | | 4.48 | 0.06 |
| 40 | 9895 - 9932 | 8758 - 8795 | 38 | | 4.16 | 0.04 |
| 41 | 381 - 397 | NA | 17 | | 4.51 | -0.02 |
| 42 | 3985 - 4015 | 3368 - 3398 | 31 | | 4.01 | -0.04 |
| 43 | 9135 - 9218 | 8127 - 8210 | 84 | | 4.26 | -0.04 |
| 44 | 8955 - 9069 | 7947 - 8061 | 115 | | 4.04 | -0.04 |
| 45 | 5835 - 6054 | 5215 - 5426 | 220 | | 3.97 | -0.11 |
| 46 | 1285 - 1308 | 752 - 775 | 24 | | 4.05 | -0.27 |
| 47 | 478 - 546 | NA | 69 | 5' Trans-activation response element (5' TAR, Roy *et al.* (1990)) | 4.16 | -0.50 |
| 48 | 6204 - 6212 | 5573 - 5581 | 9 | | 2.64 | -0.55 |
| 49 | 9258 - 9266 | 8250 - 8258 | 9 | | 3.12 | -0.55 |
| 50 | 4031 - 4058 | 3414 - 3441 | 28 | | 4.41 | -0.62 |
| 51 | 5761 - 5769 | 5141 - 5149 | 9 | | 3.22 | -0.70 |
| 52 | 3588 - 3596 | 2971 - 2979 | 9 | | 3.20 | -0.73 |
| 53 | 9661 - 9672 | NA | 12 | | 3.72 | -0.74 |
| 54 | 3646 - 3655 | 3029 - 3038 | 10 | | 3.40 | -0.76 |
| 55 | 10021 - 10030 | 8884 - 8893 | 10 | | 3.14 | -0.76 |
| 56 | 3685 - 3817 | 3068 - 3200 | 133 | | 3.29 | -0.77 |
| 57 | 4862 - 4875 | 4245 - 4258 | 14 | Central polypurine tract (CPPT, Siegfried *et al.* (2014)) | 3.65 | -0.78 |
| 58 | 6092 - 6192 | 5464 - 5561 | 101 | | 3.81 | -0.80 |
| 59 | 1994 - 2002 | NA | 9 | | 2.67 | -0.83 |
| 60 | 7856 - 7875 | NA | 20 | | 2.93 | -0.86 |
| 61 | 4134 - 4145 | 3517 - 3528 | 12 | | 3.23 | -0.86 |
| 62 | 7987 - 8000 | 7018 - 7031 | 14 | | 3.26 | -0.87 |
| 63 | 9940 - 9986 | 8803 - 8849 | 47 | | 3.09 | -0.87 |
| 64 | 7687 - 7709 | 6805 - 6821 | 23 | | 3.24 | -0.91 |
| 65 | 3396 - 3536 | 2779 - 2919 | 141 | | 3.60 | -0.98 |
| 66 | 9080 - 9096 | 8072 - 8088 | 17 | | 2.82 | -1.05 |
| 67 | 5799 - 5834 | 5179 - 5214 | 36 | | 4.06 | -1.06 |
| 68 | 5566 - 5602 | 4949 - 4985 | 37 | | 3.23 | -1.14 |
| 69 | 6727 - 6875 | NA | 149 | | 3.03 | -1.18 |
| 70 | 2234 - 2563 | 1677 - 1946 | 330 | Gag-pol frameshift (Chamorro *et al.*, 1992) | 3.68 | -1.21 |
| 71 | 6068 - 6078 | 5440 - 5450 | 11 | | 2.03 | -1.22 |
| 72 | 6646 - 6659 | 5985 - 5998 | 14 | SP stem (Siegfried *et al.* (2014)) | 3.13 | -1.23 |
| 73 | 1895 - 1901 | 1362 - 1368 | 7 | | 2.00 | -1.23 |
| 74 | 6403 - 6443 | 5757 - 5788 | 41 | | 3.05 | -1.24 |
| 75 | 3867 - 3878 | 3250 - 3261 | 12 | | 3.32 | -1.25 |
| 76 | 7715 - 7729 | NA | 15 | | 2.57 | -1.27 |
| 77 | 6547 - 6583 | 5886 - 5922 | 37 | SP stem (Siegfried *et al.* (2014)) | 3.26 | -1.27 |
| 78 | 4440 - 4463 | 3823 - 3846 | 24 | | 3.10 | -1.29 |

| Rank | Alignment position | Mapped position | Length | Name and reference | Median | z-score |
|---|---|---|---|---|---|---|
| 79 | 1231 - 1263 | 704 - 730 | 33 | | 2.93 | -1.29 |
| 80 | 4084 - 4092 | 3467 - 3475 | 9 | | 3.18 | -1.31 |
| 81 | 1954 - 1976 | 1421 - 1443 | 23 | | 3.42 | -1.34 |
| 82 | 995 - 1026 | 501 - 532 | 32 | 5' Untranslated region (5'UTR, Siegfried *et al.* (2014)) | 3.62 | -1.35 |
| 83 | 7290 - 7298 | 6417 - 6425 | 9 | | 1.74 | -1.36 |
| 84 | 3166 - 3214 | 2549 - 2597 | 49 | | 3.36 | -1.38 |
| 85 | 3624 - 3641 | 3007 - 3024 | 18 | | 2.44 | -1.38 |
| 86 | 1314 - 1394 | 781 - 861 | 81 | | 3.31 | -1.42 |
| 87 | 7395 - 7409 | 6522 - 6536 | 15 | | 1.32 | -1.52 |
| 88 | 4390 - 4403 | 3773 - 3786 | 14 | | 3.00 | -1.53 |
| 89 | 5784 - 5792 | 5164 - 5172 | 9 | | 2.19 | -1.59 |
| 90 | 2030 - 2052 | 1482 - 1498 | 23 | | 2.48 | -1.61 |
| 91 | 2110 - 2197 | 1556 - 1640 | 88 | Gag-pol frameshift (Chamorro *et al.*, 1992) | 3.73 | -1.69 |
| 92 | 4466 - 4499 | 3849 - 3882 | 34 | | 2.49 | -1.75 |
| 93 | 4938 - 4962 | 4321 - 4345 | 25 | Central polypurine tract (CPPT, Siegfried *et al.* (2014)) | 1.90 | -1.81 |
| 94 | 7742 - 7759 | 6848 - 6862 | 18 | | 2.70 | -1.84 |
| 95 | 6590 - 6600 | 5929 - 5939 | 11 | SP stem (Siegfried *et al.* (2014)) | 1.84 | -1.86 |
| 96 | 9605 - 9637 | 8475 - 8507 | 33 | | 2.21 | -1.89 |
| 97 | 8805 - 8869 | 7800 - 7864 | 65 | | 3.22 | -1.91 |
| 98 | 1933 - 1953 | 1400 - 1420 | 21 | | 1.99 | -2.02 |
| 99 | 9098 - 9117 | 8090 - 8109 | 20 | | 2.70 | -2.03 |
| 100 | 8925 - 8942 | 7920 - 7937 | 18 | | 1.97 | -2.07 |
| 101 | 6461 - 6472 | 5806 - 5817 | 12 | | 1.46 | -2.08 |
| 102 | 2986 - 3113 | 2369 - 2496 | 128 | | 3.09 | -2.11 |
| 103 | 6504 - 6522 | 5846 - 5861 | 19 | SP stem (Siegfried *et al.* (2014)) | 2.17 | -2.20 |
| 104 | 9273 - 9354 | NA | 82 | | 3.22 | -2.21 |
| 105 | 3883 - 3922 | 3266 - 3305 | 40 | | 2.65 | -2.28 |
| 106 | 9564 - 9573 | 8434 - 8443 | 10 | | 1.50 | -2.29 |
| 107 | 7880 - 7898 | 6956 - 6962 | 19 | | 2.26 | -2.32 |
| 108 | 7028 - 7258 | 6245 - 6385 | 231 | | 3.02 | -2.32 |
| 109 | 9364 - 9398 | NA | 35 | | 2.71 | -2.33 |
| 110 | 5192 - 5541 | 4575 - 4924 | 350 | | 3.69 | -2.34 |
| 111 | 6284 - 6391 | 5638 - 5745 | 108 | | 2.38 | -2.36 |
| 112 | 7783 - 7822 | 6886 - 6925 | 40 | | 2.53 | -2.42 |
| 113 | 5114 - 5133 | 4497 - 4516 | 20 | | 1.91 | -2.52 |
| 114 | 9221 - 9238 | 8213 - 8230 | 18 | | 2.39 | -2.60 |
| 115 | 6885 - 6922 | NA | 38 | | 1.99 | -2.67 |
| 116 | 8580 - 8606 | 7593 - 7607 | 27 | | 1.32 | -2.87 |
| 117 | 8622 - 8762 | 7623 - 7757 | 141 | | 2.61 | -2.92 |
| 118 | 4991 - 5072 | 4374 - 4455 | 82 | | 2.65 | -3.25 |
